# Tree diversity effects on productivity depend on mycorrhizae and life strategies in a temperate forest experiment

**DOI:** 10.1101/2022.04.13.488178

**Authors:** Peter Dietrich, Olga Ferlian, Yuanyuan Huang, Shan Luo, Julius Quosh, Nico Eisenhauer

## Abstract

Tree species are known to predominantly interact either with arbuscular mycorrhizal (AM) or ectomycorrhizal (EM) fungi. However, there is a knowledge gap whether these mycorrhizae differently influence biodiversity-ecosystem functioning (BEF) relationships and whether a combination of both can increase community productivity. In 2015, we established a tree-diversity experiment by growing tree communities with varying species-richness levels (1, 2, or 4 species), and either with AM or EM tree species, or a combination of both. We investigated basal area and annual basal area increment from 2015 to 2020 as proxy for community productivity. We found significant positive relationships between tree species richness and community productivity, which strengthened over time. Further, AM and EM tree species differently influenced productivity; however, there was no overyielding when AM and EM trees grew together. EM tree communities were characterized by low productivity in the beginning, but an increase of increment over time, and showed overall strong biodiversity effects. For AM tree communities the opposite was true. While young trees did not benefit from the presence of the other mycorrhizal type, dissimilar mechanisms underlying BEF relationships in AM and EM trees indicate that maximizing tree and mycorrhizal diversity may increase ecosystem functioning in the long run.

## Introduction

Numerous studies have shown a positive biodiversity-ecosystem functioning (BEF) relationship in terrestrial ecosystems, and that this relationship strengthens over time (Bongers et al., 2021; Cardinale et al., 2007; Fargione et al., 2007; Guerrero-Ramirez et al., 2017; Huang et al., 2018; Reich et al., 2012; Urgoiti et al., 2022). Typically, this relationship is explained by complementary resource use among plants, as well as a growing intensity of negative plant-soil interactions at low diversity and positive interactions at high diversity (Barry et al., 2019; Eisenhauer 2012). Strong evidence was found for the first two mechanisms, while research on the role of positive plant-soil interactions are, by contrast, underrepresented and often provided inconclusive results (Barry et al., 2019; Wright et al., 2017). This is surprising considering that soil-borne mutualists play a fundamental role for plants by influencing nutrient availability, pathogen pressure or competition relationships and thereby have a key impact on plant growth (Bennett et al., 2006; Tedersoo et al., 2020; Wagg et al., 2011). One of the possible reasons for inconsistent results could be that plant-mutualist relationships are not always beneficial to the plants. They can vary between beneficial and detrimental depending on factors like environmental conditions, plant age, and developmental state of the association (Johnson et al., 1997). Considering this, it is not yet fully understood whether and how plant-soil mutualist interactions contribute to the strengthening of positive BEF relationship (Wright et al., 2017).

Among soil-borne mutualists, there is one group that is ubiquitous and has an extensive influence on plants: mycorrhizal fungi (Tedersoo et al., 2020). These fungi form symbiotic relationships with almost all terrestrial plants (92% of families) and significantly influence plant productivity by enhancing nutrient acquisition, water supply and/or protection against pathogens, in exchange for photosynthetically derived carbon (Bennett et al., 2006; Smith and Read 2008; Wang and Qiu 2006). There are two main types of mycorrhizal symbiosis, arbuscular mycorrhizae (AM), which are common in many different plant families, and ectomycorrhizae (EM), which occur only in certain families (mainly tree and shrub families; Wang and Qiu (2006)). Both differ considerably in their structure and cause dissimilar effects on plant growth: AM are more critical for phosphorous uptake, while EM are mainly involved in nitrogen mineralisation and uptake (Holste et al., 2017; Read 1991). Moreover, EM associations are known to have a competitive advantage over AM, for example, due to saprotrophic activity and the development of a denser hyphal network (Jones et al., 1998; Malloch et al., 1980; Read and Perez-Moreno 2003), but are also more resource costly for plants and thus only beneficial under specific conditions (Rygiewicz and Andersen 1994).

Almost all tree species are predominantly associated with either AM or EM fungi (Ferlian et al., 2021), therefore are classified as AM or EM tree species (Wang and Qiu 2006). Tree associations with specific mycorrhizal types may be related to their nutrient acquisition strategies and functional traits (Averill et al., 2019; Cornelissen et al., 2001). Specifically, AM tree species mainly rely on inorganic nutrient resources and therefore request fast nutrient cycling and acquisitive traits, while EM tree species can benefit from saprotrophic activity of EM fungi and thus are associated with conservative traits in slow-nutrient-cycling ecosystems. Those differences between AM and EM tress may have influenced their life strategies: most AM tree species are fast-growing, while most EM tree species are slow-growing species (Myking et al., 2013; Petrokas et al., 2020). However, this does not always match, for example, the EM associated *Betula pendula* Roth is a classic pioneer species that can grow very fast (Petrokas et al., 2020).

Given that AM and EM associations are fundamentally different (fungi as well as trees), it is feasible to assume that all partners benefit from growing together (Ferlian et al., 2018). In such AM+EM tree communities, resource use complementarity may be highest due to an efficient exploitation of the available belowground and aboveground resources, i.e., soil nutrients and light via differences in growth strategies and leaf trait expressions (Ferlian et al., 2018; Grossman et al., 2017; Wagg et al., 2011). To test this assumption, we established a tree-diversity experiment (MyDiv) by growing tree communities with varying species richness (1, 2, or 4), and either with AM or EM tree species, or a combination of both (50:50). This experimental design has several advantages and can answer still-open questions:

- Most knowledge about BEF relationships are based on findings in grasslands, but mechanical understanding behind BEF relationship in forest ecosystem are still rare. It is necessary to test the existing knowledge in forest ecosystems to be able to formulate general conclusions.
- It is still not well understood how AM and EM trees contribute to BEF relationships and how this change over time. This is not only important for BEF research, but also has relevance for forest management and the selection of tree species for plantations and reforestation.
- Previous biodiversity experiments have shown that diverse plant communities produce more biomass and have higher diversity/abundance of soil-borne mutualists, but it is not clear who influenced whom (Barry et al., 2019) under which conditions (e.g., along a plant diversity gradient). The use of AM and EM trees as model organisms can help to answer this question by a more direct manipulation of the functional diversity of mycorrhizae. This allows for a closer look at the role of soil mutualists in BEF relationships and to disentangle causes from consequences.

Here, we used plot-level basal area and annual basal area increment as proxy for productivity (Bongers et al., 2020; Hutchison et al., 2018; Schnabel et al., 2019), and tested how tree species richness and mycorrhizal type (only AM/EM trees, or both AM and EM trees) interactively influenced productivity during the establishment phase of the MyDiv experiment (2015-2020).

Moreover, we tested whether potential positive diversity effects were caused by complementarity among tree species or by high productivity of few species in mixtures (= selection effects), using the additive partitioning method of Loreau and Hector (2001). Thereby, we hypothesized that H1) tree species-rich communities are more productive than tree species-poor communities and that this difference increases over time. This is explainable by differences in biodiversity effects (complementarity vs. selection effects), with initially high selection effects and later high complementarity effects in high-diversity mixtures, and generally low biodiversity effects in low-diversity mixtures.

H2) communities containing both AM and EM tree species show a higher productivity than communities with only AM or EM tree species. Differences among AM, EM and AM+EM tree communities become greater with time.

H3) there is an interaction between species richness and mycorrhizal type, i.e., AM+EM tree communities with four tree species have highest productivity, while EM only communities have lowest productivity. This is explainable by different impacts of AM and EM species on biodiversity effects caused by differences in life strategies (fast vs. slow).

## Materials and methods

### Study site and experimental design

The MyDiv experiment is a long-term tree diversity experiment located near Halle (Saxony-Anhalt, Germany; 51°23’ N,11°53’ E) at the Bad Lauchstädt Experimental Research Station of the Helmholtz Centre for Environmental Research - UFZ (Ferlian et al., 2018). The elevation is 115 m a.s.l. and the climate is continental. Mean annual air temperature for 2005-2014 (decade before the experiment was established) was 10.0°C, and mean annual precipitation was 535.3 mm, both recorded with an automatic meteorological station near the experimental site (Meteorological data of Bad Lauchstädt, Helmholtz Centre for Environmental Research - UFZ, Department of Soil System Science. https://www.ufz.de/index.php?de=39439). The soil is a Haplic Chernozem developed from Loess with a pH range between 6.6 and 7.4 (Altermann et al., 2005; Ferlian et al., 2018). Moreover, soil is rich in nitrogen, while phosphorous is more limited (Supporting Information S1 Fig. S1; Ferlian *et al*. (2018)). Before the experiment was established in March 2015, the site had been used for agriculture until 2012 and as a grassland from 2013-2015. The study site was divided into two blocks due to a gradient of abiotic and biotic parameters measured before the establishment of the experiment (Ferlian et al., 2018).

The experiment consists of 80 11 × 11 m plots that are covered with a water-permeable weed tarp (minimization of weed interference), with a core area of 8 × 8 m (64 m^2^). In March 2015, 140 two-year-old tree seedlings (nursery plants, bare-rooted, 50–80 cm height, no mycorrhiza inoculation) were planted in a 1 × 1 m grid in each plot (64 trees in the core area). The species pool included ten tree species (Table 1). The selection of species was made with a focus on similarity, i.e., all species are deciduous angiosperms, are native to Germany, and are adapted to site conditions including the ability to tolerate high light exposure (in the beginning) and shade (at later stages; Ferlian et al. 2018). The only difference is that the tree species interact with different types of mycorrhiza, whereby five of them predominantly associate with AM and the other five with EM fungi (Table 1). This enabled us to test for pure mycorrhizae type effects, although we cannot exclude that the selection of tree species also has some influence on the results. There was no direct control of mycorrhizal fungal association; and the treatment was established through assignment of tree species to dominant mycorrhizae based on literature review (Ferlian et al., 2018). Assessments of mycorrhization in 2019 in the MyDiv experiment confirmed these assignments (Ferlian et al., 2021). Tree species-richness levels range from one to four species (1, 2, 4), whereas species richness was crossed with mycorrhizal type (i.e., two- and four-species mixtures contain either only AM or EM tree species, or both AM and EM tree species (50:50)), while there was no mixed mycorrhizal type in monocultures. Per species, there are two monoculture replicates, one in each block (N=20). Furthermore, ten replicates per species richness level and mycorrhizal type (AM, EM, AM+EM) were established, distributed over the two blocks (N_two-species_=30; N_four-species_=30). More information about the design of the MyDiv experiment can be found in Ferlian et al., (2018).

**Table 1.** Overview of tree species used in the MyDiv experiment (scientific and common name), their mycorrhizal associations (AM=arbuscular mycorrhizae, EM=ectomycorrhizae), life strategies, maximum age (in years), maximum tree height (in m), tree height in 2020 (in m; five years after the establishment of the experiment), and increment over time, i.e., increase or decrease of basal area increment during study period (2015-2020). Information about mycorrhizal association and max. tree height were derived from Ferlian et al., (2018), max. age from Brzeziecki and Kienast (1994); life strategies were determined by cluster analysis; tree height in 2020 was calculated as mean of tree height of all tree individuals in 2020, and increment over time as the slope of simple linear regression between basal area increment (mean of both monocultures per species) and year (2016-2020).

| Tree species | Common name | Mycor-rhizal association | Life strategy | Max. age (year) | Max. tree height (m) | Tree height in 2020 (m) | Increment (m<sup>2</sup> m<sup>-2</sup> year<sup>-1</sup>) |
| --- | --- | --- | --- | --- | --- | --- | --- |
| *Acer pseudoplatanus* L. | Sycamore, maple | AM | Fast | 300 | 20 | 5.12 | -0.06 |
| *Aesculus hippocastanum* L. | Horse chestnut | AM | Fast | 200 | 20 | 2.95 | 0.05 |
| *Fraxinus excelsior* L. | Ash | AM | Slow | 300 | 25 | 4.37 | 0.03 |
| *Prunus avium* (L.) L. | Bird/wild cherry | AM | Fast | 150 | 20 | 5.18 | -0.20 |
| *Sorbus aucuparia* L. | Rowan, rowanberry | AM | Fast | 60 | 10 | 4.32 | -0.01 |
| *Betula pendula* Roth | Silver birch | EM | Fast | 120 | 17.5 | 6.18 | -0.36 |
| *Carpinus betulus* L. | Hornbeam | EM | Slow | 250 | 15 | 3.91 | 0.12 |
| *Fagus sylvatica* L. | Copper beech | EM | Slow | 300 | 25 | 2.39 | 0.21 |
| *Quercus petraea* (Matt.) Liebl. | Sessile oak | EM | Slow | 500 | 27.5 | 2.36 | 0.21 |
| *Tilia platyphyllos* Scop. | Large-leaved lime | EM | Slow | 400 | 27.5 | 4.08 | 0.29 |

### Data collection and calculations

Tree diameter was measured five cm above the ground (= basal diameter; d_0_; m) for all 64 trees in the core area of the plots for basal area calculation, with a diameter tape, annually in December/January 2015 – 2020. Vitality of the trees was assessed at the same time. Trees were classified as dead when the phloem was clearly dead (brown instead of green), or the whole tree was missing (for example, by trees falling over). From 2015 to 2018, dead trees were replaced by new individuals that had grown next to the experimental site since March 2015 (same age, same growth habit, similar environmental conditions). Replacement was low: 3.3% (168 out of 5120 trees) in the first year and less then 1.5% in all subsequent years (for more species- and richness-level-specific information about replacement and number of missing trees, see Supporting Information S1 Table S1 and S2).

With tree diameter, we calculated standing basal area (m^2^ m^-2^) in each year and annual basal area increment (m^2^ m^-2^ year^-1^) of the tree communities as proxy for productivity (= ecosystem functioning). Dead trees that were missing in 2019 and 2020 were treated as zero values, whereas the percentage was low (2019: 0.18% missing trees; 2020: 1.95% missing trees; Supporting Information S1 Table S1). To account for differences among years, which were caused, inter alia, by differences in weather conditions (Supporting Information S1 Fig. S2), we detrended the calculated variables (basal area, annual basal area increment) by dividing each value per plot by the averaged value of all plots per year (Dietrich et al., 2020).

Finally, we calculated net biodiversity effects (NEs), selection effects (SEs), and complementarity effects (CEs) for basal area (absolute and detrended data) by using the additive partitioning method of Loreau and Hector (2001), separately for each year and tree community. These biodiversity effects depend on relative yields of species, which was calculated using monoculture values as denominator. Thereby, we used monoculture values of block 1 for biodiversity effect calculations of block 1 mixtures, and monoculture values of block 2 for block 2 mixtures. We skipped biodiversity effects calculation based on annual basal area increment, because some communities (especially monocultures) had a very low annual increment, which resulted in unrealistically high biodiversity effect values. For species-specific SEs and CEs (based on basal area), we used the following equations

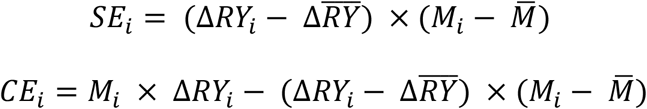

where *ΔRY*_*i*_ is the deviation from expected relative yield of species *i* in the mixture (RY_observed_-RY_expected_), and *M*_*i*_ is the yield of species *i* in monoculture (Huang et al., 2018). Net biodiversity effects (NEs) were calculated by adding SE and CE per plot and year. RY_observed_ (hereafter referred to as RY) of AM and EM trees in AM+EM tree communities was further used to study how trees perform in presence of the other mycorrhizal type (i.e., in AM+EM communities).

To test whether positive diversity effects are caused by the different life strategies of the AM and EM trees we used the fast-slow approach (Reich 2014). To do so, we grouped tree species of the MyDiv experiment into fast and slow species, related to nutrient acquisition, growth strategy, and life span. The grouping can be understood as different types of life strategies (fast vs. slow), similar to the concept of different strategies of trees during succession (pioneers vs. climax species). We clustered tree species based on the following variables: maximum age, maximum growth height, wood density, specific leaf area, leaf C/N ratio, and basal area increment of monoculture communities over time (Huston and Smith 1987; Reich 2014; Westoby and Wright 2006). All variables were derived from Ferlian et al., (2018); except maximum age, which came from Brzeziecki and Kienast (1994), and increment, which was calculated as the slope of simple linear regression between basal area increment (mean of both monocultures per species) and year (2016-2020). Clustering revealed that five out of ten species have a fast life strategy, and other species have a slow life strategy (Supporting Information S2 Fig. S1), based on Reich (2014), and Huston and Smith (1987), and in line with Reuter et al., (2021). Slow species were characterized by high growth height and maximum age, low specific leaf area (SLA), and an increase of basal area increment over time; while fast species showed the opposite (Supporting Information S2 Table S1; Fig. S2). Four out of five slow species were EM tree species (+ AM tree species *F. excelsior*), and four out of five fast species were AM tree species (+ EM tree species *B. pendula*; Table 1; Supporting Information S2 Fig. S1). We then grouped the tree communities into three community-strategy types: (1) tree communities containing more than 50% fast species (i.e., 75% or 100%) were considered as fast communities (31 communities), (2) tree communities with more than 50% slow species as slow communities (31 communities), and (3) tree communities containing 50% fast and 50% slow species as fast+slow communities (18 communities).

### Statistical analysis

To test whether tree communities performed differently depending on tree species richness, mycorrhizal type and whether this changed over time, linear mixed-effects models were fitted with basal area, annual basal area increment (absolute and detrended data, respectively), and biodiversity effects (NEs, SEs, CEs) as response variables, respectively. We used block, plot, and the interaction of block and year as random effects and started with a null model with these random effects only. Then, we added the fixed effects tree species richness (1, 2, 4; log-transformed), mycorrhizal type (AM, EM, AM+EM), year (2015/16-2020), and all possible interactions in a sequential order. Prior to analyses, biodiversity effects (NEs, SEs, CEs) were square-root-transformed with sign reconstruction (sign(y)=|y|) to improve the normality of residuals (Loreau and Hector 2001).

To test whether missing trees influenced community productivity, we added density of living trees per plot as a covariate before the other fixed effects in separate models with absolute and detrended basal area as response variables. Further, we performed sensitivity analyses to test whether specific tree species played a significant role in community performance. We excluded all plots containing a specific tree species from the dataset and analyzed data as described above (for basal area). To test the impact of community strategies (fast, slow, fast+slow) on tree productivity, we used the same models as described above, but replaced mycorrhizal type (AM, EM, AM+EM) with community strategy. Differences among mycorrhizal types or community strategies in a specific year were tested with mixed-effects models with block as random and mycorrhizal type/community strategy as fixed effect, followed by Tukey’s HSD test. All models were fitted with maximum likelihood (ML), and likelihood ratio tests were used to decide on the significance of the fixed effects.

To test whether biodiversity effects per year and group (mycorrhizal type/community strategy/species richness) were significantly greater than zero, we used one-tailed t-tests. The strength of the relationships between response variables and year, and response variables and tree species richness were tested with simple linear regression analysis using the slope of the regression line (positive slope indicate increase, negative slope indicate decrease, increase in slope over years indicate strengthening and decrease of slope weakening of relationship). All calculations and statistical analyses were done in R (version 3.6.1, R Development Core Team, http://www.R-project.org), including the package *lme4* (Bates et al., 2015) for mixed-effects model analysis, *multcomp* (Hothorn et al., 2016) for Tukey HSD test, and *vegan* (Oksanen et al., 2007) for PCA (Supporting Information S2 Fig. S2).

## Results

Overall, we found similar results for absolute and detrended data (Table 2 and 3), indicating that the impacts of tree species richness, mycorrhizal type, and community strategy on community productivity was not strongly affected by yearly fluctuations (although the effect size may have been influenced). Note that we calculated stem biomass and their annual increment as another proxy of productivity, the results of which were similar to that of basal area (see Supporting Information S3 for biomass calculation and results).

**Table 2.** Summary of mixed-effects model analyses testing the effects of log-transformed tree species richness (1, 2, 4; Sr), mycorrhizal type (AM, EM, AM+EM; Myc), year (2015/16-2020), and their interactions (upper part of the table), and log-transformed tree species richness, community strategy (fast, slow, fast+slow; CS), year, and their interactions (lower part of the table) on basal area and their annual increment (absolute and detrended values). Degrees of freedom (DF), Chi^2^- and P-values (P) are shown. Significant effects (P<0.05) are given in bold and marginally significant effects (P<0.1) in italics. Note that detrended values were calculated by dividing absolute value per plot by the averaged value of all plots per year, to account for differences among years due to climatic fluctuations.

|  |  | Basal area |  |  |  | Annual increment |  |  |  |
| --- | --- | --- | --- | --- | --- | --- | --- | --- | --- |
|  |  | absolute |  | detrended |  | absolute |  | detrended |  |
|  | Df | Chi<sup>2</sup> | P | Chi<sup>2</sup> | P | Chi<sup>2</sup> | P | Chi<sup>2</sup> | P |
| Tree species richness (Sr) | 1 | 5.98 | **0.014** | 3.55 | *0.060* | 7.71 | **0.005** | 9.35 | **0.002** |
| Mycorrhizal type (Myc) | 2 | 19.36 | **<0.001** | 28.89 | **<0.001** | 4.79 | *0.091* | 2.95 | 0.229 |
| Year | 1 | 43.31 | **<0.001** | 0.00 | 0.995 | 3.21 | *0.073* | 0.00 | 0.998 |
| Sr x Myc | 2 | 1.58 | 0.453 | 0.93 | 0.627 | 2.28 | 0.320 | 2.27 | 0.322 |
| Sr x Year | 1 | 29.81 | **<0.001** | 5.69 | **0.017** | 2.89 | *0.089* | 1.18 | 0.278 |
| Myc x Year | 2 | 13.29 | **0.001** | 110.44 | **<0.001** | 24.63 | **<0.001** | 18.99 | **<0.001** |
| Sr x Myc x Year | 2 | 7.36 | **0.025** | 5.56 | *0.062* | 0.51 | 0.774 | 0.27 | 0.875 |
|  | Df | Chi<sup>2</sup> | P | Chi<sup>2</sup> | P | Chi<sup>2</sup> | P | Chi<sup>2</sup> | P |
| Tree species richness (Sr) | 1 | 5.98 | **0.014** | 3.55 | *0.060* | 7.71 | **0.005** | 9.35 | **0.002** |
| Community strategy (CS) | 2 | 21.55 | **<0.001** | 37.22 | **<0.001** | 1.62 | 0.445 | 0.50 | 0.778 |
| Year | 1 | 43.36 | **<0.001** | <0.01 | 0.995 | 3.21 | *0.073* | <0.01 | 0.998 |
| Sr x CS | 2 | 4.61 | *0.100* | 6.08 | **0.048** | 1.11 | 0.573 | 0.80 | 0.670 |
| Sr x Year | 1 | 29.81 | **<0.001** | 5.69 | **0.017** | 2.89 | *0.089* | 1.18 | 0.278 |
| LS x Year | 2 | 4.64 | *0.098* | 201.69 | **<0.001** | 55.43 | **<0.001** | 48.24 | **<0.001** |
| Sr x CS x Year | 2 | 3.96 | 0.138 | 11.43 | **0.003** | 10.17 | **0.006** | 8.20 | **0.017** |

**Table 3.** Summary of mixed-effects model analyses testing the effects of log-transformed tree species richness (2, 4; Sr), mycorrhizal type (AM, EM, AM+EM; Myc), year (2015-2020), and their interactions on net diversity effects (NEs), selection effects (SEs), and complementarity effects (CEs) based on absolute and detrended basal area values. Shown are degrees of freedom (DF), Chi^2^- and P-values (P). Significant effects (P < 0.05) are given in bold and marginally significant effects (P < 0.1) in italics. Note that detrended basal area values were calculated by dividing absolute value per plot by the averaged value of all plots per year, to account for differences among years, e.g., due to climatic fluctuations. Furthermore, we did not show biodiversity effects for annual increment, because some communities (especially monocultures) had a very low annual increment, which resulted in unrealistically high biodiversity effect values.

| Basal area (absolute) |  |  |  |  |  |  |  |
| --- | --- | --- | --- | --- | --- | --- | --- |
|  | NEs |  |  | SEs |  | CEs |  |
|  | Df | Chi<sup>2</sup> | P | Chi<sup>2</sup> | P | Chi<sup>2</sup> | P |
| Tree species richness (Sr) | 1 | 2.67 | 0.102 | 6.85 | **0.009** | 0.09 | 0.768 |
| Mycorrhiza type (Myc) | 2 | 4.81 | *0.090* | 10.23 | **0.006** | 0.45 | 0.797 |
| Year | 1 | 12.36 | **<0.001** | 2.28 | 0.131 | 4.16 | **0.041** |
| Sr x Myc | 2 | 2.41 | 0.300 | 1.21 | 0.545 | 1.22 | 0.542 |
| Sr x Year | 1 | 12.48 | **<0.001** | 2.13 | 0.145 | 3.48 | *0.062* |
| Myc x Year | 2 | 9.27 | **0.010** | 1.51 | 0.471 | 1.27 | 0.531 |
| Sr x Myc x Year | 2 | 5.60 | *0.061* | 4.61 | *0.100* | 4.30 | 0.117 |
| Basal area (detrended) |  |  |  |  |  |  |  |
|  | NEs |  |  | SEs |  | CEs |  |
|  | Df | Chi<sup>2</sup> | P | Chi<sup>2</sup> | P | Chi<sup>2</sup> | P |
| Tree species richness (Sr) | 1 | 1.36 | 0.244 | 6.56 | **0.010** | 0.00 | 0.960 |
| Mycorrhiza type (Myc) | 2 | 3.90 | 0.142 | 12.14 | **0.002** | 0.63 | 0.728 |
| Year | 1 | 2.15 | 0.143 | 0.99 | 0.320 | 0.13 | 0.715 |
| Sr x Myc | 2 | 1.63 | 0.442 | 1.11 | 0.575 | 0.81 | 0.666 |
| Sr x Year | 1 | 11.80 | **0.001** | 1.48 | 0.223 | 6.38 | **0.012** |
| Myc x Year | 2 | 4.58 | 0.101 | 0.11 | 0.946 | 0.52 | 0.772 |
| Sr x Myc x Year | 2 | 3.57 | 0.168 | 4.43 | 0.109 | 3.17 | 0.205 |

### Hypothesis 1: Diversity-productivity relationships increase over time

Basal area and basal area increment significantly increased with tree species richness (Table 2; Fig. 1). Moreover, we found a significant interaction between tree species richness and linear year on basal area, even if we controlled for differences among years (detrended values; Table 2), which indicates that positive diversity effects can develop and increase over time (Fig. 1; Supporting Information S1 Table S3). Those patterns remained the same after accounting for the effects of tree density (Supporting Information S1 Table S4) and the identity of tree species (Supporting Information S1 Table S5).

**Figure 1.**
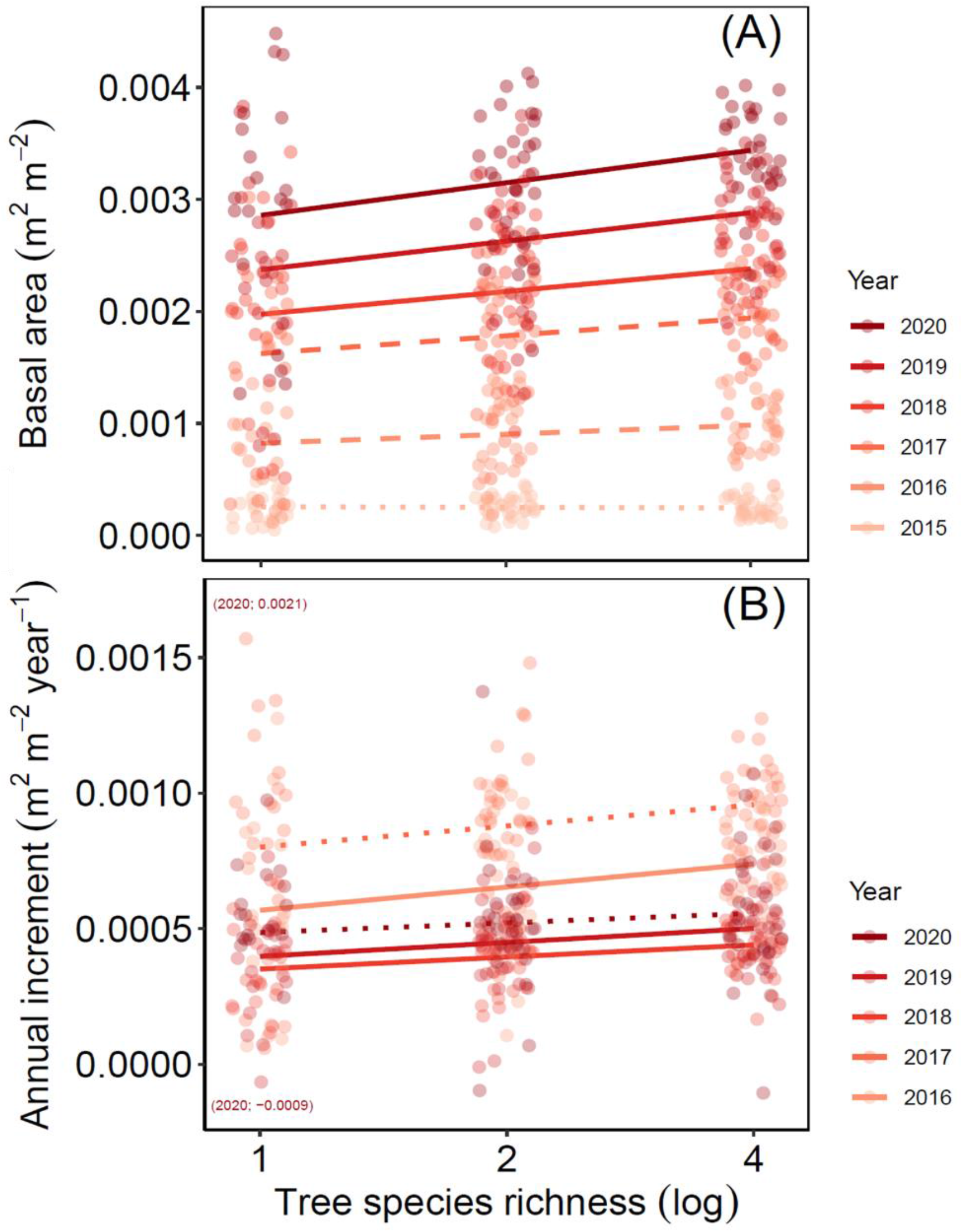
Basal area (A) and its annual increment (B) as function of log-transformed tree species richness from 2015/16 to 2020. Each dot represents a tree community and colors indicate different years. Regression lines are based on mixed-effects models (predicted means), while solid lines indicate significant relationships (P<0.05), dashed lines marginally significant relationships (0.1>P>0.05), and dotted lines non-significant relationships (P>0.1). Note that two data points were excluded from the figure, because they have either very high positive or negative values. These data points are indicated as text in parentheses (year; value).

We found positive NEs, which increased over the years (Table 3; Fig. 2a). This was initially caused by a steady increase of SEs from 2015 to 2019 (Fig. 2a). Complementarity effects (CEs) also increased in the beginning (CEs higher than SEs in 2016), remained stable from 2017-2018 (CEs lower than SEs), and then continued to increase (CEs higher than SEs; Fig. 2a). From 2019 to 2020, SEs strongly decreased, thus NEs were mainly attributed by CEs in 2020 (Fig. 2a). All biodiversity effects were significantly higher than zero (except for NEs and SEs in 2015, and SEs in 2020; Fig. 2a; Supporting Information S1 Table S6). Overall, this pattern was found in two- and four-tree species communities in similar way, while there were also differences between these communities: we found higher SEs and a stronger increase in CEs in four-than in two-species communities, which results in higher NEs (and CEs) in four-than in two-species communities in 2020 (Table 3; Fig. 2b, c; Supporting Information S1 Table S7).

**Figure 2.**
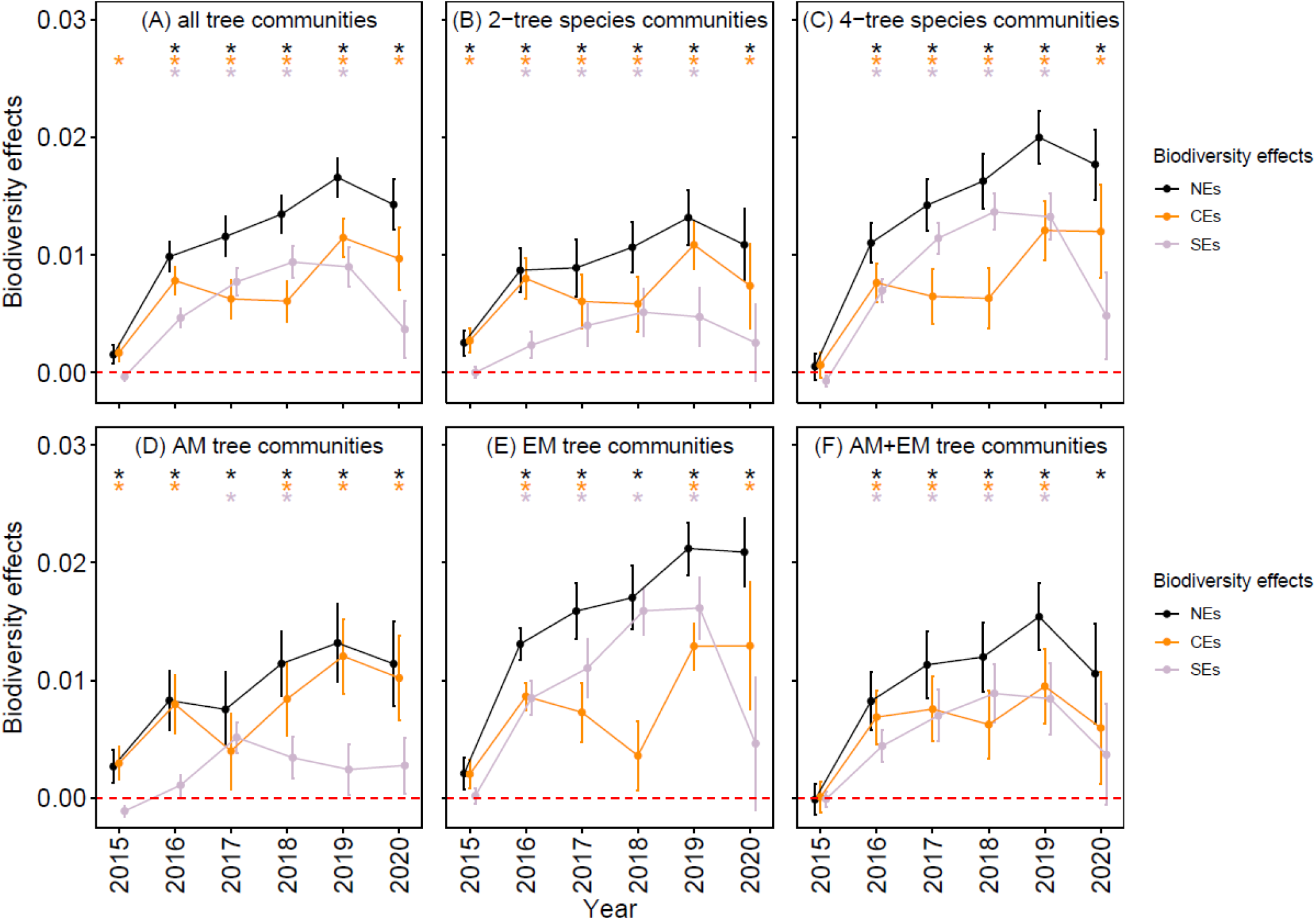
Changes in net diversity effects (NEs), complementarity effects (CEs), and selection effects (SEs; different colors) over time for all mixture tree communities (A), separately for two- and four species communities (B, C), and for communities containing AM (C), EM (D), or both AM and EM tree species (E). Dots represent mean values (± standard error) per year, and asterisks indicate whether the biodiversity effects were significantly greater than zero (red dotted line = zero line; detailed t-test results can be found in Supporting Information S1 Table S6, S7, and S11). The y-axes are square root–transformed to reflect the quadratic nature of biodiversity effects (Loreau and Hector 2001).

### Hypothesis 2: Tree communities assembled with different mycorrhizal types differ in productivity over time

After eight months’ growth (in December 2015), basal area differed among communities assembled by different mycorrhizal types (P<0.001, Table 1), with highest basal area for AM tree communities and lowest for EM tree communities, and this pattern was maintained during the entire study period and in all species richness levels (Fig. 3a). However, differences in basal area of AM, EM, and AM+EM tree communities decreased over time (Supporting Information S1 Table S8). This was due to changes in basal area increment over the years: increment was highest in AM tree communities, and lowest in EM tree communities after eight months’ growth (Fig. 4a; Supporting Information S1 Table S9), but decreased in AM tree communities (P<0.001), remained stable in AM+EM tree communities (P=0.335), and increased in EM tree communities (P=0.014) over time, leading to the highest annual increment in EM tree communities in 2020 (Fig. 4a).

**Figure 3.**
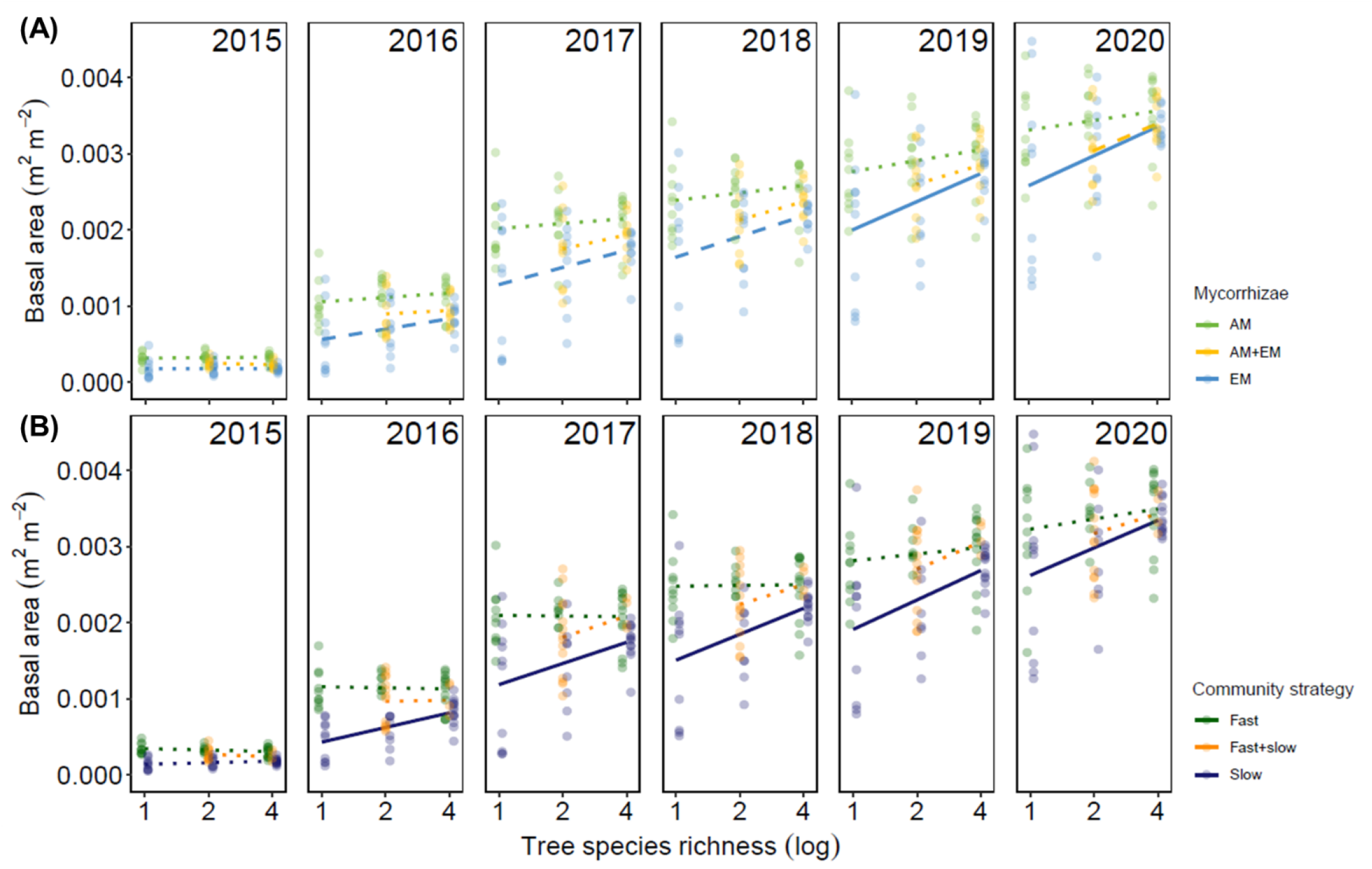
Basal area as function of log-transformed tree species richness for 2015 to 2020, for communities containing AM, EM, or both AM and EM tree species (A), and for communities containing fast, slow, or both fast and slow tree species (B). Each dot represents a tree community, and colors indicate different mycorrhizal types and community strategies. Regression lines are based on mixed-effects models (predicted means), while solid lines indicate significant relationships (P<0.05), dashed lines marginally significant relationships (0.1>P>0.05), and dotted lines non-significant relationships (P>0.1).

**Figure 4.**
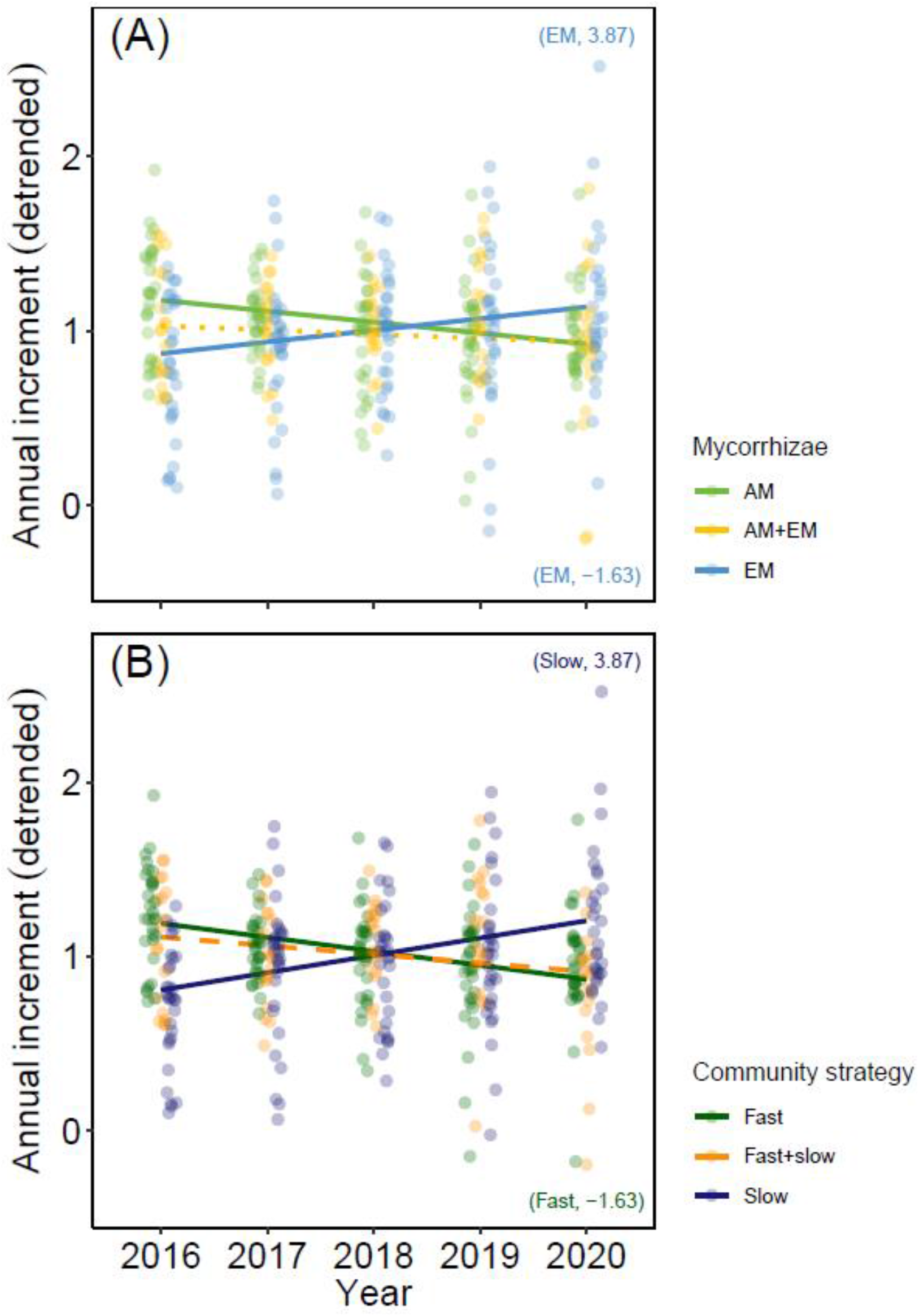
Annual increment of basal area over time from 2016 to 2020, for communities containing AM, EM, or both AM and EM tree species (A), and for communities containing fast, slow, or both fast and slow tree species (B). Annual increment was detrended, i.e., each value per plot was divided by the averaged value of all plots per year, to account for differences among years (e.g., climatic fluctuations). Each dot represents a tree community, and colors indicate different mycorrhizal types and community strategies. Regression lines are based on mixed-effects models (predicted means), while solid lines indicate significant relationships (P<0.05), dashed lines marginally significant relationships (0.1>P>0.05), and dotted lines non-significant relationships (P>0.1). Note that two data points were excluded from the figure, because they have either very high positive or negative values. These data points are indicated as text in parentheses (mycorrhizal type/community strategy; value).

Comparison of RY of AM and EM trees in AM+EM communities showed that there was no difference in the first two years of the experiment, while from 2017 to 2020 RY of AM trees was higher than RY of EM trees (mean of RY_AM_ > 0.5; mean of RY_EM_ < 0.5; Supporting Information S1 Fig. S3). Moreover, RY of AM trees increased over time, while RY of EM trees decreased, in AM+EM communities (Supporting Information S1 Fig. S3).

### Hypothesis 3: Mycorrhizal types differentially influence biodiversity effects due to differences in life strategies

We found a significant three-way interaction effect (species richness x mycorrhizal type x year) on basal area (Table 2), which can be explained by differences in the relationships between species richness and basal area in AM, EM, and AM+EM tree communities over the years (Fig. 3). At the beginning of the experiment (2015), there was no significant relationship between species richness and basal area in all communities. In 2016-2020, AM tree communities also showed no significant relationship (Fig. 3a; Supporting Information S1 Table S10); whereas EM tree communities had higher basal area in four-species communities than in monocultures (2016-2018: marginally significant; 2019-2020: significant; Supporting Information S1 Table S10). In AM+EM tree communities, basal area tended to increase with increasing species richness in 2020 (Fig. 3a; Supporting Information S1 Table S10). For all three-types of communities, the slope of species richness-basal area relationship increased over the years (except that the slope in AM tree communities decreased from 2019 to 2020; Supporting Information S1 Table S10).

We found significant differences in biodiversity effects among communities with different mycorrhizal types: EM tree communities showed the steepest increase of NEs, resulting in highest NEs in 2020 (Table 3; Fig. 2d-f). These NEs were mainly caused by SEs until 2019. In 2020, NEs were mainly caused by CEs due to the decrease of SEs (Fig. 2e; Supporting Information S1 Table S11). In AM tree communities, NEs were primarily caused by CEs in the whole period, while SEs were absent (2015, 2016, 2020) or weak (2017-2019; Fig. 2d; Supporting Information S1 Table S11). In AM+EM tree communities, CEs and SEs contributed equally to NEs (Fig. 2f; Supporting Information S1 Table S11).

Species-specific analyses revealed that a few species strongly contributed to changes in SEs and CEs over the years: one of the most influential species was the EM tree species *B. pendula*, which showed the highest basal area in mixtures, especially in four-species mixtures (Supporting Information S1 Fig. S4). Consequently, this species had highes SE values among all species from 2015 to 2018 in four-species communities, while SEs were lower in two-species communities (Supporting Information S1 Fig. S5, S6). In 2020, *B. pendula* showed a strong decrease in SEs, which caused the overall loss in SEs in 2020 (Supporting Information S1 Fig. S5). Other species, which showed a strong increase in SEs, mainly in four-species communities, were the EM species *F. sylvatica* and *Q. petraea* and the AM species *P. avium* (Supporting Information S1 Fig. S6). For species-specific CEs, again *B. pendula* had the most influential effect: CEs of this species showed the strongest increase, and this increase tended to be higher in four-than in two-species communities (Supporting Information S1 Fig. S6). The AM tree species *A. pseudoplatanus* and *P. avium* also showed an increase of CEs over the years, while the EM tree species *C. betulus, F. sylvatica*, and *Q. petraea*, and the AM tree species *A. hippocastanum* showed a decrease in CEs over time (Supporting Information S1 Fig. S5).

There were significant differences in basal area among the three types of communities based on community strategies (Table 2). Overall, fast communities had highest, and slow communities lowest basal area (Fig. 3b). One exception were four-species communities, where fast+slow communities had equal or slightly higher basal area than fast communities (significant interaction of tree species richness and community strategy; Table 2; Fig. 3b). Next to this result, replacing community strategy with mycorrhizal type in our model did not provide new insights (since most fast tree species are AM associated and most slow tree species are EM associated). Therefore, remaining results of community strategy analysis (e.g., for basal area increment and biodiversity effects) can be found in Table 2, Figure 4b, and Supporting Information S2.

## Discussion

### Hypothesis 1: Diversity-productivity relationships increase over time

We found a significant positive effect of tree species richness on basal area, and the effect steadily increased over the years. This is in line with previous findings in grasslands and forests showing that biodiversity-ecosystem functioning relationships become steeper over time (Bongers et al., 2021; Guerrero-Ramirez et al., 2017; Huang et al., 2018; Reich et al., 2012; Schnabel et al., 2019). Notably, we found a significant effect of species richness on annual basal area increment already in the second year of the experiment (2016) and then in almost all subsequent years. This shows that tree communities can benefit from higher species richness during the very early establishment phase. This can be explained by both selection and complementarity effects in the beginning of our study. Highly-productive pioneer species, such as *B. pendula* or *P. avium*, caused strong selection effects, and can function as shelter species in mixtures by reducing high light levels and extreme temperature changes, providing nutrients (high leaf N and P concentration, fast litter decomposition), and protection from wind, supporting neighbouring tree species during establishment (Cornelissen et al., 2001; Löf et al., 2014; Stark et al., 2015; Wright et al., 2017). Moreover, differences in belowground nutrient use and/or aboveground growth forms (canopy packing) among species can increase resource use efficiency, complementarity effects, and thus productivity in mixtures (Bongers et al., 2021; Morin et al., 2011; Reuter et al., 2021; Williams et al., 2021). As pioneer species are fast-growing, selection effects increased more strongly in the first years of the experiment (especially in four-species mixtures), but increasing biotic interactions and changing competitive interactions altered this trajectory (Bongers et al., 2021). As a result, selection effects decreased after four years in our experiment, while complementarity effects continued to increase. This is in line with previous findings showing that selection effects play a major role at the beginning, but are replaced by complementarity effects after some time (Fargione et al., 2007; Jing et al., 2021; Lasky et al., 2014). Overall, our results support the hypothesis that co-occurring species in high-diverse temperate forests differ in niches and competitive abilities, which leads to strong biodiversity effects and higher productivity in mixtures compared to their monospecific counterparts (Bongers et al., 2021; Huang et al., 2018; Morin et al., 2011; Urgoiti et al., 2022). Our results extend this knowledge by showing that both complementarity and selection effects contribute to biodiversity effects on tree growth in young tree stands.

### Hypothesis 2: Tree communities assembled with different mycorrhizal types differ in productivity over time

In contrast to our second hypothesis, we did not find overyielding when both mycorrhizal types were present in the community. Consistently, AM and EM trees did not show complementary use of resources, which is in line with a previous study in the MyDiv experiment (Reuter et al., 2021). In fact, AM tree communities had the highest basal area in all diversity levels and over the whole study period, which is consistent with a recent study in a subtropical forest (Ma et al., 2021). The high productivity of AM tree communities could be due to two reasons: on the one hand, AM tree species are known to have a fast growth strategy (Cornelissen et al., 2001; Ferlian et al., 2018), and several species are pioneers at juvenile stages, e.g., *S. aucuparia* (Myking et al., 2013) and *P. avium* (Petrokas et al., 2020); on the other hand, the soil of the experimental field is rich in nitrogen, while phosphorous is more limited (Ferlian et al., 2018), which favours AM associations more than EM associations (Holste et al., 2017; Read 1991). This can also explain why AM trees dominated in the AM+EM communities, i.e., AM trees had increasing RY over time, while the opposite was true for EM trees, which is in line with previous studies (Bennett et al., 2017; Chen et al., 2019; Mao et al., 2019). The different growth strategies may also explain why AM communities decreased in increment while EM communities increased increment over time. Previous work has shown that fast-growing species lose and slow-growing species increase productivity, partly due to differences in nutrient economy and interactions with soil mutualists (Dietrich et al., 2020).

### Hypothesis 3: Mycorrhizal types differentially influence biodiversity effects due to differences in life strategies

We found a significant positive diversity-productivity relationship in EM tree communities, whereas no such relationships were found in AM tree communities. The positive diversity effect in EM tree communities can be explained by two consecutive processes: First, selection effects dominated from 2016 to 2019, when the EM species *B. pendula* was the most-productive species causing strong selection effects. However, also two other EM species, *F. sylvatica* and *Q. petraea*, caused high selection effects due to their very low productivity, across all species richness levels. The phenomenon that species cause positive selection effects, when they were low-productive in monocultures *and* mixtures, were also found in previous work (Huang et al., 2018). The fast growth of *B. pendula*, with highest productivity across all species, and slow growth of *F. sylvatica* and *Q. petraea*, with lowest productivity across all species, indicates that growth strategies of EM tree species cover a broad spectrum (from very fast to very slow species), while in AM communities, the growth strategies of the species were relatively similar in our experiment (Supporting Information S1 Figure S7: change in basal area increment over time strongly differed among EM species, while in AM species, there was a relative similar change in increment over time). The fundamentally different growth strategies of EM tree species can be an explanation for stronger biodiversity effects found in our study and is in line with previous assumptions (Brzeziecki and Kienast 1994; Ferlian et al., 2018; Niklaus et al., 2017). The loss of many *B. pendula* individuals in monoculture in 2020 and thus a loss of productivity can explain why the selection effect strongly decreased from 2019 to 2020.

Second, we found an increase in complementarity effects over time in EM tree communities, which is explainable by a disproportionate increase of complementarity effects of *B. pendula* compared to other tree species. *B. pendula* had an intermediate productivity in monoculture in our experiment, but strongly increased its productivity with tree species richness representing the most-productive species in four-species mixtures. Consequently, *B. pendula* was responsible for the positive complementarity effect in EM tree communities, while three out of four other EM tree species showed a weak negative complementarity effect in 2020. With our data in hand, we can only speculate why *B. pendula* is much more productive in mixture. The most likely explanation is that *B. pendula* benefited from other EM trees growing more slowly, so that there was more light available, while in monocultures, birch trees were in strong competition with each other (Jucker et al., 2014; Williams et al., 2021). A further explanation could be plant-soil feedback effects (PSFs), which were shown to be main drivers of positive BEF relationships in grasslands and forests (Bennett et al., 2017; Eisenhauer et al., 2019; Guerrero-Ramírez et al., 2019; Thakur et al., 2021). For example, it is possible that high mortality in *B. pendula* monocultures in 2020 was caused by pathogen accumulation (= negative PSFs); however, it is also possible that harsh weather conditions, such as summer drought, led to greater mortality (Schnabel et al., 2021; Schnabel et al., 2019; Steckel et al., 2020). Thus, future studies should address in more detail how PSFs (especially negative PSFs), extreme weather events and their combined effect contribute to BEF relationships in the MyDiv experiment.

In AM tree communities, there was no significant BEF relationship; nevertheless, we found strengthening complementarity effects. Currently, we can only speculate regarding the underlying drivers (e.g., plant-soil feedback effects). Presumably there was no significant BEF relationship because of highly productive AM monocultures reflecting that AM trees are often pioneer species.

An explanation for different biodiversity effects in AM and EM communities are differences in life strategies. This is supported by our cluster analyses showing that most fast tree species were AM associated and most slow tree species were EM associated (Averill et al., 2019; Cornelissen et al., 2001), and that we therefore did not find strong differences between analyses using mycorrhizal type (AM, EM, AM+EM) and using community strategy (fast, slow, fast+slow) as fixed effect. Nevertheless, we gained one important new insight: tree communities containing fast and slow species had similar or slightly higher productivity than fast or slow communities alone. This result supports the hypothesis that an increase in the number of species pursuing different strategies (e.g., life strategies) can increase ecosystem functioning (Hao et al., 2020; Niklaus et al., 2017; Teste et al., 2017). We found this effect only in four-species mixtures, indicating that two species may not make full use of the available niches, and that both high tree species richness and high variability in life strategies are needed to sustain a high ecosystem functioning in forest ecosystems (Bongers et al., 2021; Niklaus et al., 2017). However, it is important to note that our community strategy analysis is very basic. Future work should consider site-specific above- and belowground plant traits and their plasticity across different tree diversity levels (Eisenhauer 2022) as well as how different mycorrhizal types (AM, EM, AM+EM) influence trait expression and their link to productivity.

## Conclusions

Our study shows that tree species richness is essential to maintain high ecosystem functioning, and that tree diversity effects increase over time. These findings provide empirical evidence that species-rich plantations can be more productive than monocultures, whereby the choice of tree species is important. Even though our experimental plots are relatively small (in comparison to the size of “real” forests), our results can serve as a basis for further investigations on larger scale with a more application-related focus (i.e., forest management and reforestation). More precisely, we found that AM and EM tree communities differ strongly in their productivity in terms of temporal development and tree diversity effects. Although we did not directly manipulate mycorrhizal fungi in the MyDiv experiment, a recent study confirmed that AM trees were significantly higher colonized by AM fungi, while EM trees were significantly higher colonized by EM fungi in the MyDiv experiment (Ferlian et al., 2021). Therefore, we are convinced that the effects we found were caused by specific fungus-tree interactions and that we were able to successfully manipulate the functional diversity of mycorrhizae. Nevertheless, we did not find overyielding when both mycorrhizal types were present in the community after five years of growing. We assume that, shortly after planting the trees in a nitrogen-rich soil, aboveground factors, such as light intensity and space competition, may play a more important role for community productivity than complementarity in soil resource use. At later growth stages, however, when nutrients become more limited, the combination of AM and EM trees may have a greater impact on ecosystem functioning. Therefore, long-term observations are needed to fully understand the role of plant-soil mutualist interactions for strengthening BEF relationships.

## Acknowledgments

We thank Ulrich Pruschitzki for help setting up the experiment and maintaining the field site in the first years. This work was supported by the European Research Council (ERC) under the European Union’s Horizon 2020 research and innovation program (grant agreement no. 677232), and by the German Research Foundation (DFG) in the frame of the Gottfried Wilhelm Leibniz Prize (Ei 862/29-1). Further support came from the German Centre for Integrative Biodiversity Research (iDiv) Halle-Jena-Leipzig, funded by the German Research Foundation (FZT 118). SL was supported by a Humboldt Research Fellowship.

## Notes

**Data availability** The data that support the findings of this study are openly available in BExIS at http://doi.org/[doi], reference number [reference number].

### Competing Interest Statement

The authors have declared no competing interest.

