## Supplement material S1 for "Tree diversity effects on productivity depend on mycorrhizae and life strategies in a temperate forest experiment"

**Table S1** Overview of dead trees that were replaced (2015-2018) or missing (2019; 2020) in the respective years: total number, number per species richness level, and number per tree species. Numbers in brackets indicate the mortality rate in relation to the total number of planted trees.

| | 2015 | 2016 | 2017 | 2018 | 2019 | 2020 |
|---|---|---|---|---|---|---|
| Total | 168 (3.3%) | 51 (1.0%) | 77 (1.5%) | 72 (1.4%) | 9 (0.2%) | 100 (2.0%) |
| Monocultures | 44 | 19 | 40 | 22 | 2 | 40 |
| Two-species plots | 61 | 17 | 27 | 26 | 4 | 28 |
| Four-species plots | 63 | 15 | 10 | 24 | 3 | 32 |
| *Acer pseudoplatanus* L. | 2 | 9 | 7 | 10 | 0 | 5 |
| *Aesculus hippocastanum* L. | 7 | 0 | 5 | 0 | 0 | 0 |
| *Betula pendula* Roth | 1 | 0 | 0 | 0 | 0 | 56 |
| *Carpinus betulus* L. | 4 | 3 | 2 | 0 | 0 | 1 |
| *Fagus sylvatica* L. | 67 | 5 | 6 | 9 | 6 | 19 |
| *Fraxinus excelsior* L. | 1 | 0 | 2 | 26 | 0 | 3 |
| *Prunus avium* (L.) L. | 1 | 0 | 0 | 2 | 0 | 0 |
| *Quercus petraea* (Matt.) Liebl. | 77 | 25 | 46 | 22 | 3 | 12 |
| *Sorbus aucuparia* L. | 5 | 1 | 1 | 0 | 0 | 0 |
| *Tilia platyphyllos* Scop. | 3 | 8 | 8 | 3 | 0 | 4 |

**Table S2** Overview of dead individuals of *Betula pendula*, *Fagus sylvatica*, and *Quercus petraea* that were replaced (2015-2018) or missing (2019; 2020), per year and species richness level (1-spec- = one tree species; 2-spec. = two tree species; 4-spec. = four tree species).

| | *Betula pendula* | | | *Fagus sylvatica* | | | *Quercus petraea* | | |
|---|---|---|---|---|---|---|---|---|---|
| | 1-spec. | 2-spec. | 4-spec. | 1-spec. | 2-spec. | 4-spec. | 1-spec. | 2-spec. | 4-spec. |
| 2015 | 0 | 0 | 1 | 19 | 24 | 24 | 16 | 28 | 34 |
| 2016 | 0 | 0 | 0 | 0 | 4 | 1 | 8 | 6 | 11 |
| 2017 | 0 | 0 | 0 | 0 | 2 | 4 | 31 | 15 | 0 |
| 2018 | 0 | 0 | 0 | 0 | 3 | 6 | 2 | 9 | 11 |
| 2019 | 0 | 0 | 0 | 0 | 4 | 2 | 2 | 0 | 1 |
| 2020 | 20 | 17 | 10 | 0 | 8 | 11 | 4 | 2 | 6 |

**Table S3** Summary of simple linear regression analyses between basal area and log-transformed tree species richness, and annual increment of basal area and log-transformed tree species richness in the years 2015-2020, respectively. Shown are degrees of freedom (DF), F- and P-values as well as slope of the regression line. Significant influences are given in bold and marginally significant influences in italics (P-values).

| | Basal area ~ species richness | | | | Annual increment ~ species richness | | | |
|---|---|---|---|---|---|---|---|---|
| | DF | F | P | slope | DF | F | P | slope |
| 2015 | 1 | <0.01 | 0.934 | $-1.7\times10^{-6}$ | - | - | - | - |
| 2016 | 1 | 3.25 | *0.075* | $1.3\times10^{-4}$ | 1 | 5.92 | **0.017** | $1.2\times10^{-4}$ |
| 2017 | 1 | 3.61 | *0.061* | $2.1\times10^{-4}$ | 1 | 2.31 | 0.132 | $8.6\times10^{-5}$ |
| 2018 | 1 | 5.53 | **0.021** | $2.7\times10^{-4}$ | 1 | 5.04 | **0.028** | $0.6\times10^{-4}$ |
| 2019 | 1 | 8.07 | **0.006** | $3.6\times10^{-4}$ | 1 | 7.14 | **0.009** | $0.9\times10^{-4}$ |
| 2020 | 1 | 6.91 | **0.010** | $3.8\times10^{-4}$ | 1 | <0.01 | 0.980 | $-1.7\times10^{-6}$ |

**Table S4** Summary of mixed-effects model analyses testing the effects of density of living trees within plot (max. 64), log-transformed tree species richness (1, 2, 4; Sr), mycorrhizal type (AM, EM, AM+EM; Myc), year (2015-2020) and their interaction on basal area (absolute and detrended values). Shown are degrees of freedom (DF), $Chi^2$, and P-values (P). Significant influences are given in bold and marginally significant influences in italics. Note that detrended values were calculated by dividing absolute value per plot by the averaged value of all plots per year, to account for differences among years due to climatic fluctuations.

| | | Basal area with density | | | | Basal area | | | |
|---|---|---|---|---|---|---|---|---|---|
| | | absolute | | detrended | | absolute | | detrended | |
| | Df | $Chi^2$ | P | $Chi^2$ | P | $Chi^2$ | P | $Chi^2$ | P |
| Density | 1 | 39.81 | **<0.001** | 23.28 | **<0.001** | - | - | - | - |
| Tree species richness (Sr) | 1 | 5.66 | **0.017** | 3.36 | *0.067* | 5.98 | **0.014** | 3.55 | *0.060* |
| Mycorrhiza type (Myc) | 2 | 17.85 | **<0.001** | 27.37 | **<0.001** | 19.36 | **<0.001** | 28.89 | **<0.001** |
| Year | 1 | 44.61 | **<0.001** | 0.67 | 0.411 | 43.31 | **<0.001** | 0.00 | 0.995 |
| Sr x Myc | 2 | 1.27 | 0.530 | 0.75 | 0.688 | 1.58 | 0.453 | 0.93 | 0.627 |
| Sr x Year | 1 | 28.75 | **<0.001** | 4.75 | **0.029** | 29.81 | **<0.001** | 5.69 | **0.017** |
| Myc x Year | 2 | 8.80 | **0.012** | 137.46 | **<0.001** | 13.29 | **0.001** | 110.44 | **<0.001** |
| Sr x Myc x Year | 2 | 5.71 | *0.058* | 5.66 | *0.059* | 7.36 | **0.025** | 5.56 | *0.062* |

**Table S5** Summary of sensitivity analyses (mixed-effects models) testing the effects of log-transformed tree species richness (1, 2, 4; Sr), mycorrhizal type (AM, EM, AM+EM; Myc), year (2015-2020) and their interaction on basal area, when plots containing specific tree species were excluded. Shown are degrees of freedom (DF), Chi$^2$, and P-values (P). Significant influences are given in bold and marginally significant influences in italics.

| | | Basal area | | | | | | | | | |
|---|---|---|---|---|---|---|---|---|---|---|---|
| | | without plots with *A. pseudoplatanus* | | without plots with *A. hippocastanum* | | without plots with *B. pendula* | | without plots with *C. betulus* | | without plots with *F. sylvatica* | |
| | Df | Chi$^2$ | P | Chi$^2$ | P | Chi$^2$ | P | Chi$^2$ | P | Chi$^2$ | P |
| Species richness (Sr) | 1 | 2.30 | 0.130 | 2.94 | *0.086* | 5.17 | **0.023** | 5.43 | **0.020** | 4.01 | **0.045** |
| Mycorrhiza type (Myc) | 2 | 10.43 | **0.005** | 13.23 | **0.001** | 23.75 | **<0.001** | 13.63 | **0.001** | 8.11 | **0.017** |
| Year | 1 | 45.14 | **<0.001** | 42.74 | **<0.001** | 43.55 | **<0.001** | 41.57 | **<0.001** | 40.84 | **<0.001** |
| Sr x Myc | 2 | 1.88 | 0.390 | 1.32 | 0.516 | 0.88 | 0.645 | 1.80 | 0.406 | 0.51 | 0.774 |
| Sr x Year | 1 | 16.65 | **<0.001** | 20.88 | **<0.001** | 18.97 | **<0.001** | 20.71 | **<0.001** | 11.72 | **0.001** |
| Myc x Year | 2 | 7.91 | **0.019** | 9.06 | **0.011** | 13.70 | **0.001** | 17.36 | **<0.001** | 4.32 | 0.115 |
| Sr x Myc x Year | 2 | 8.20 | **0.017** | 2.54 | 0.280 | 5.25 | *0.072* | 5.64 | *0.060* | 3.38 | 0.184 |

| | | Basal area | | | | | | | | | |
|---|---|---|---|---|---|---|---|---|---|---|---|
| | | without plots with *F. excelsior* | | without plots with *P. avium* | | without plots with *Q. petraea* | | without plots with *S. aucuparia* | | without plots with *T. platyphyllos* | |
| | Df | Chi$^2$ | P | Chi$^2$ | P | Chi$^2$ | P | Chi$^2$ | P | Chi$^2$ | P |
| Species richness (Sr) | 1 | 3.21 | *0.073* | 5.55 | **0.018** | 5.05 | **0.025** | 3.44 | *0.064* | 5.77 | **0.016** |
| Mycorrhiza type (Myc) | 2 | 18.28 | **<0.001** | 7.90 | **0.019** | 8.80 | **0.012** | 23.22 | **<0.001** | 25.53 | **<0.001** |
| Year | 1 | 44.93 | **<0.001** | 45.89 | **<0.001** | 39.20 | **<0.001** | 45.07 | **<0.001** | 44.14 | **<0.001** |
| Sr x Myc | 2 | 0.23 | 0.890 | 1.30 | 0.522 | 0.41 | 0.814 | 0.06 | 0.973 | 1.74 | 0.419 |
| Sr x Year | 1 | 21.92 | **<0.001** | 25.55 | **<0.001** | 13.04 | **<0.001** | 21.37 | **<0.001** | 30.90 | **<0.001** |
| Myc x Year | 2 | 17.56 | **<0.001** | 3.10 | 0.213 | 2.92 | 0.233 | 31.02 | **<0.001** | 51.81 | **<0.001** |
| Sr x Myc x Year | 2 | 1.52 | 0.467 | 6.05 | **0.048** | 1.65 | 0.438 | 0.95 | 0.622 | 10.29 | **0.006** |

**Table S6** Summary of one-tailed t-test analyses (values higher than zero) for net diversity effects (NEs), complementarity effects (CEs) and selection effect (SEs) of basal area (absolute and detrended values) from 2015-2020. Shown are degrees of freedom (DF), t- and P-values. Significant influences are given in bold and marginally significant influences in italics (P-values).

| | | Basal area (absolute) | | | | | | Basal area (detrended) | | | | | |
|---|---|---|---|---|---|---|---|---|---|---|---|---|---|
| | | NEs | | CEs | | SEs | | NEs | | CEs | | SEs | |
| | DF | t | P | t | P | t | P | t | P | t | P | t | P |
| 2015 | 59 | 1.47 | *0.073* | 1.71 | **0.047** | -0.66 | 0.744 | 1.47 | *0.073* | 1.71 | **0.046** | -0.67 | 0.745 |
| 2016 | 59 | 7.02 | **<0.001** | 5.46 | **<0.001** | 4.91 | **<0.001** | 7.09 | **<0.001** | 5.46 | **<0.001** | 4.91 | **<0.001** |
| 2017 | 59 | 6.23 | **<0.001** | 3.69 | **<0.001** | 5.34 | **<0.001** | 6.24 | **<0.001** | 3.69 | **<0.001** | 5.34 | **<0.001** |
| 2018 | 59 | 7.08 | **<0.001** | 3.34 | **<0.001** | 6.12 | **<0.001** | 7.08 | **<0.001** | 3.40 | **<0.001** | 6.12 | **<0.001** |
| 2019 | 59 | 8.38 | **<0.001** | 5.83 | **<0.001** | 5.14 | **<0.001** | 8.38 | **<0.001** | 5.83 | **<0.001** | 5.14 | **<0.001** |
| 2020 | 59 | 6.02 | **<0.001** | 3.37 | **<0.001** | 0.32 | 0.376 | 6.02 | **<0.001** | 3.37 | **<0.001** | 0.32 | 0.376 |

**Table S7** Summary of one-tailed t-test analyses (values higher than zero) for net diversity effect (NEs), complementarity effects (CEs) and selection effect (SEs) of basal area from 2015-2020 for AM, EM and AM+EM communities. Shown are degrees of freedom (DF), t- and P-values. Significant influences are given in bold and marginally significant influences in italics (P-values).

| | | 2-species communities | | | | | | 4-species communities | | | | | |
|---|---|---|---|---|---|---|---|---|---|---|---|---|---|
| | | NEs | | CEs | | SEs | | NEs | | CEs | | SEs | |
| | DF | t | P | t | P | t | P | t | P | t | P | t | P |
| 2015 | 29 | 1.20 | **0.028** | 2.10 | **0.022** | 0.12 | 0.454 | 0.05 | 0.478 | 0.29 | 0.386 | -1.16 | 0.873 |
| 2016 | 29 | 4.19 | **<0.001** | 3.64 | **<0.001** | 2.08 | **0.023** | 5.76 | **<0.001** | 4.08 | **<0.001** | 4.89 | **<0.001** |
| 2017 | 29 | 3.24 | **0.001** | 2.33 | **0.014** | 2.04 | **0.025** | 5.70 | **<0.001** | 2.85 | **0.004** | 5.94 | **<0.001** |
| 2018 | 29 | 4.07 | **<0.001** | 2.49 | **0.009** | 2.66 | **0.006** | 6.15 | **<0.001** | 2.32 | **0.014** | 6.33 | **<0.001** |
| 2019 | 29 | 5.37 | **<0.001** | 4.37 | **<0.001** | 2.07 | **0.023** | 6.91 | **<0.001** | 4.01 | **<0.001** | 5.44 | **<0.001** |
| 2020 | 29 | 3.07 | **0.002** | 1.71 | **0.049** | 0.04 | 0.483 | 5.55 | **<0.001** | 2.99 | **0.003** | 0.41 | 0.342 |

**Table S8** Summary of mixed-effects model and Tukey HSD analyses testing the difference in basal area among mycorrhizal types (AM, EM, AM+EM) for the years 2015-2020, respectively. Shown are degrees of freedom (DF), $Chi^2$, and P-values for mixed-effect model analysis, and z- and P-values for Tukey HSD analysis. Significant influences are given in bold and marginally significant influences in italics (P-values).

| | Mixed-effects model | | | Tukey HSD | | | | | |
|---|---|---|---|---|---|---|---|---|---|
| | | | | EM vs. AM | | AM+EM vs. AM | | AM+EM vs. EM | |
| | DF | $Chi^2$ | P | z | P | z | P | z | P |
| 2015 | 2 | 48.86 | **<0.001** | -8.12 | **<0.001** | -4.13 | **<0.001** | -3.13 | **0.005** |
| 2016 | 2 | 27.77 | **<0.001** | -5.70 | **<0.001** | -2.38 | **0.045** | -2.72 | **0.018** |
| 2017 | 2 | 19.57 | **<0.001** | -4.64 | **<0.001** | -1.74 | 0.191 | -2.41 | **0.042** |
| 2018 | 2 | 18.21 | **<0.001** | -4.45 | **<0.001** | -1.63 | 0.234 | -2.36 | **0.048** |
| 2019 | 2 | 12.84 | **0.002** | -3.64 | **<0.001** | -1.11 | 0.509 | -2.15 | *0.080* |
| 2020 | 2 | 7.50 | **0.024** | -2.77 | **0.016** | -1.17 | 0.469 | -1.30 | 0.393 |

**Table S9** Summary of mixed-effects model and Tukey HSD analyses testing the difference in detrended basal area increment among mycorrhizal types (AM, EM, AM+EM) for the years 2015-2020, respectively. Shown degrees of freedom (DF), Chi$^2$, and P-values for mixed-effect model analysis, and z- and P-values for Tukey HSD analysis. Significant influences are given in bold and marginally significant influences in italics (P-values). Note that detrended values were calculated by dividing absolute value per plot by the averaged value of all plots per year, to account for differences among years due to climatic fluctuations.

| | Mixed-effects model | | | Tukey HSD | | | | | |
|---|---|---|---|---|---|---|---|---|---|
| | | | | EM vs. AM | | AM+EM vs. AM | | AM+EM vs. EM | |
| | DF | Chi$^2$ | P | z | P | z | P | z | P |
| 2016 | 2 | 18.81 | **<0.001** | -4.54 | **<0.001** | -1.67 | 0.215 | -2.38 | **0.045** |
| 2017 | 2 | 5.70 | *0.058* | -2.34 | *0.051* | -0.61 | 0.812 | -1.48 | 0.302 |
| 2018 | 2 | 0.04 | 0.980 | 0.09 | 0.996 | 0.20 | 0.979 | -0.12 | 0.993 |
| 2019 | 2 | 1.20 | 0.549 | 0.74 | 0.737 | 1.04 | 0.551 | -0.37 | 0.925 |
| 2020 | 2 | 1.65 | 0.438 | 0.89 | 0.646 | -0.42 | 0.908 | 1.22 | 0.444 |

**Table S10** Summary of simple linear regression analyses between basal area and log-transformed tree species richness of AM, EM and AM+EM communities in the years 2015-2020, respectively. Shown are degrees of freedom (DF), F- and P-values as well as slope of the regression line. Significant influences are given in bold and marginally significant influences in italics (P-values).

| | | AM communities | | | EM communities | | | AM+EM communities | | |
|---|---|---|---|---|---|---|---|---|---|---|
| | DF | F | P | slope | F | P | slope | F | P | slope |
| 2015 | 1 | 0.19 | 0.668 | <0.0001 | <0.01 | 0.987 | <0.0001 | 0.75 | 0.399 | <0.0001 |
| 2016 | 1 | 1.02 | 0.321 | 0.00008 | 3.46 | *0.073* | 0.00020 | 0.15 | 0.699 | 0.00007 |
| 2017 | 1 | 0.55 | 0.463 | 0.00010 | 2.90 | *0.100* | 0.00032 | 1.06 | 0.317 | 0.00026 |
| 2018 | 1 | 1.09 | 0.305 | 0.00014 | 3.87 | *0.059* | 0.00039 | 2.09 | 0.166 | 0.00034 |
| 2019 | 1 | 1.57 | 0.220 | 0.00021 | 5.56 | **0.026** | 0.00053 | 1.71 | 0.207 | 0.00036 |
| 2020 | 1 | 1.10 | 0.303 | 0.00018 | 4.48 | **0.043** | 0.00056 | 3.70 | *0.071* | 0.00052 |

**Table S11** Summary of one-tailed t-test analyses (values higher than zero) for net diversity effect (NEs), complementarity effects (CEs) and selection effect (SEs) of basal area from 2015-2020 for AM, EM and AM+EM communities. Shown are degrees of freedom (DF), t- and P-values. Significant influences are given in bold and marginally significant influences in italics (P-values).

| | | | | | | | |
|---|---|---|---|---|---|---|---|
| | | | | AM communities | | | |
| | | NEs | | CEs | | SEs | |
| | DF | t | P | t | P | t | P |
| 2015 | 19 | 1.93 | **0.034** | 2.16 | **0.022** | -1.91 | 0.964 |
| 2016 | 19 | 3.09 | **0.003** | 2.93 | **0.004** | 1.16 | 0.130 |
| 2017 | 19 | 2.29 | **0.017** | 1.47 | *0.079* | 3.59 | **<0.001** |
| 2018 | 19 | 3.07 | **0.003** | 2.34 | **0.015** | 1.99 | **0.031** |
| 2019 | 19 | 3.36 | **0.002** | 2.92 | **0.004** | 1.48 | *0.078* |
| 2020 | 19 | 2.70 | **0.007** | 2.27 | **0.018** | 1.33 | *0.100* |
| | | | | EM communities | | | |
| | | NEs | | CEs | | SEs | |
| | DF | t | P | t | P | t | P |
| 2015 | 19 | 0.64 | 0.266 | 0.67 | 0.256 | 0.35 | 0.366 |
| 2016 | 19 | 6.33 | **<0.001** | 5.24 | **<0.001** | 4.95 | **<0.001** |
| 2017 | 19 | 5.49 | **<0.001** | 2.95 | **0.004** | 4.35 | **<0.001** |
| 2018 | 19 | 5.41 | **<0.001** | 1.61 | *0.062* | 5.94 | **<0.001** |
| 2019 | 19 | 6.77 | **<0.001** | 4.94 | **<0.001** | 4.52 | **<0.001** |
| 2020 | 19 | 6.09 | **<0.001** | 2.51 | **0.011** | -0.11 | 0.542 |
| | | | | AM+EM communities | | | |
| | | NEs | | CEs | | SEs | |
| | DF | t | P | t | P | t | P |
| 2015 | 19 | -0.06 | 0.524 | -0.06 | 0.524 | -0.03 | 0.510 |
| 2016 | 19 | 3.36 | **0.002** | 2.68 | **0.007** | 2.70 | **0.007** |
| 2017 | 19 | 3.30 | **0.002** | 2.15 | **0.022** | 2.58 | **0.009** |
| 2018 | 19 | 3.91 | **<0.001** | 1.83 | **0.041** | 3.48 | **0.001** |
| 2019 | 19 | 4.92 | **<0.001** | 3.05 | **0.003** | 3.04 | **0.003** |
| 2020 | 19 | 2.38 | **0.014** | 1.40 | *0.088* | 0.45 | 0.330 |

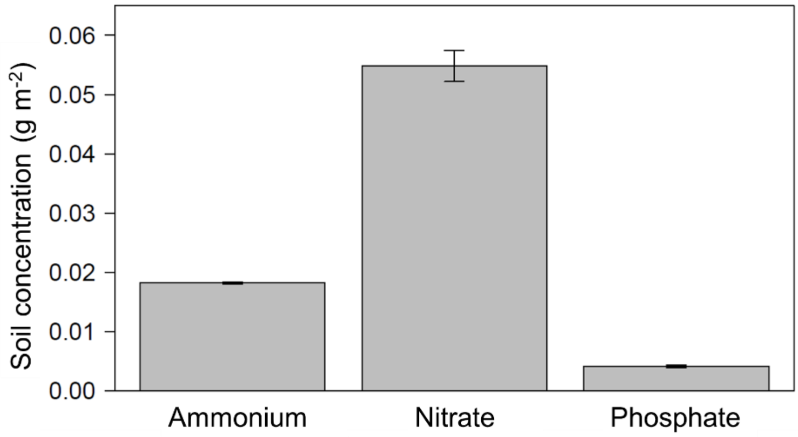


**Figure S1** Concentration (g $m^{-2}$) of ammonium ($NH_4$), nitrate ($NO_3$) and phosphate ($PO_4$) in soil of the study site. Shown are means (±standard error) of study plots.

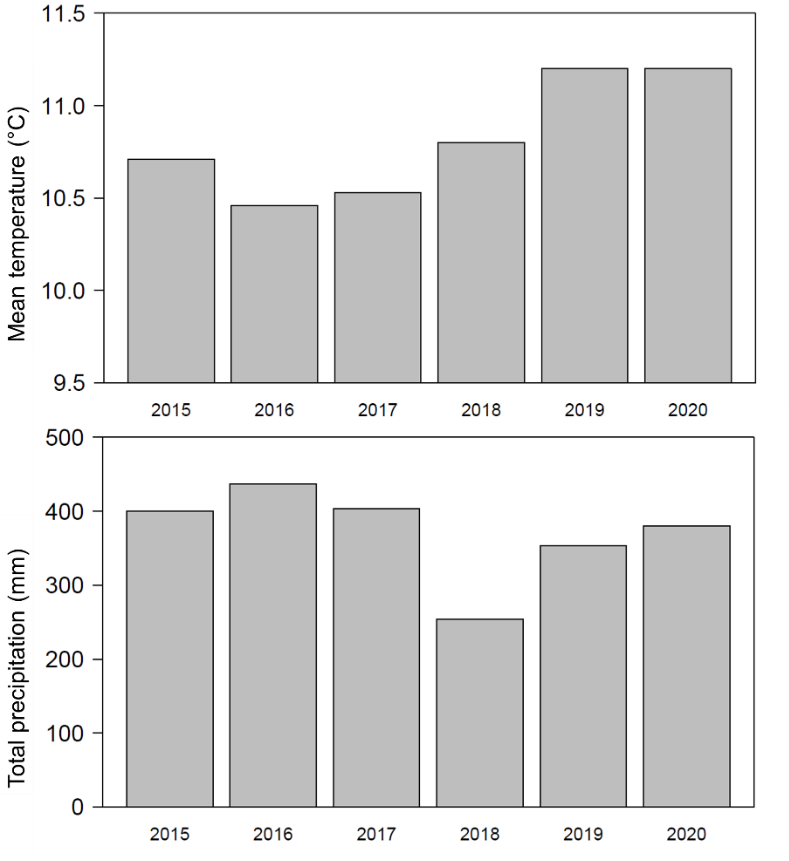


**Figure S2** Mean annual air temperature and total precipitation from 2015 to 2020. Data from 2015 to 2017 were achieved from the Helmholtz Centre for Environmental Research (UFZ) weather station (https://www.ufz.de/index.php?de=39439), and from 2018 to 2020 from the German Meteorological Service (DWD) weather station in Bad Lauchstädt (https://www.ufz.de/index.php?de=46100).

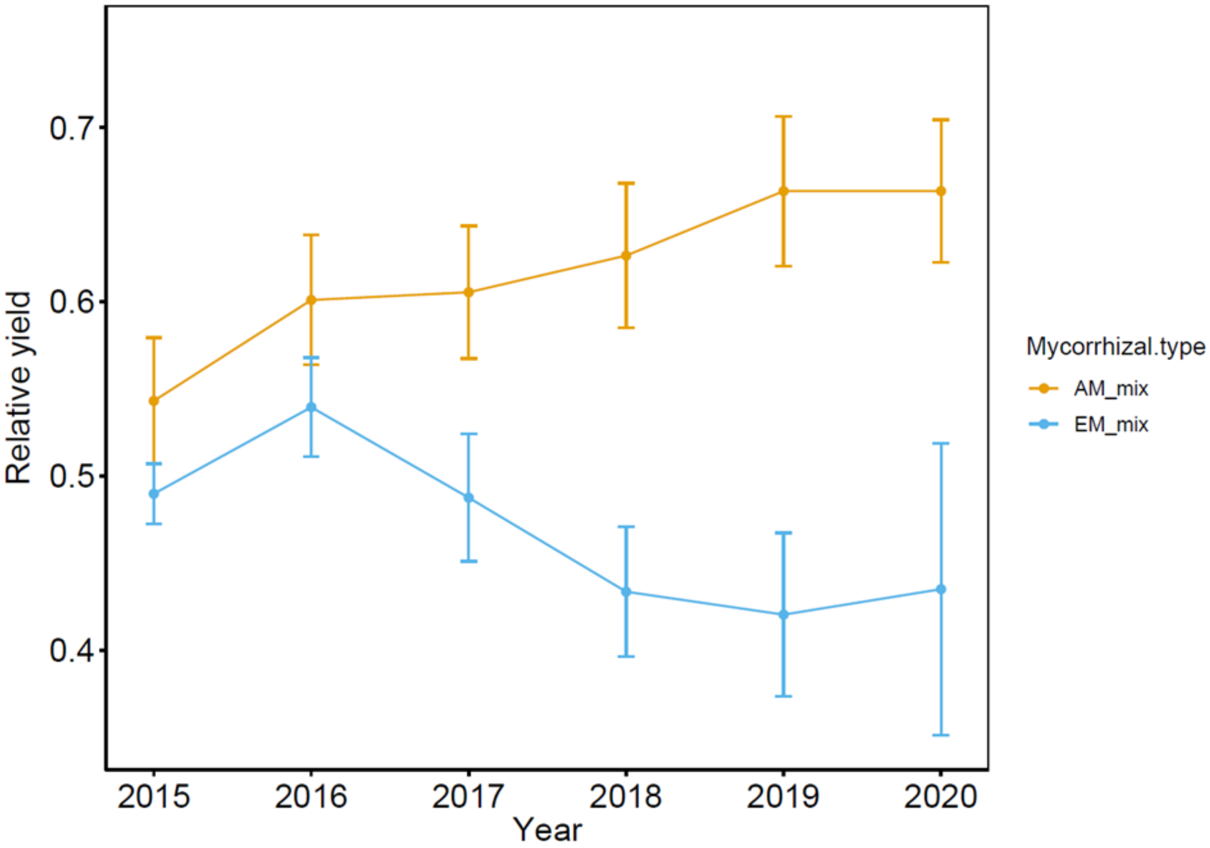


**Figure S3** Changes in relative yield (RY) of AM and EM trees in AM+EM communities. Dots represent mean values (± standard error) per year.

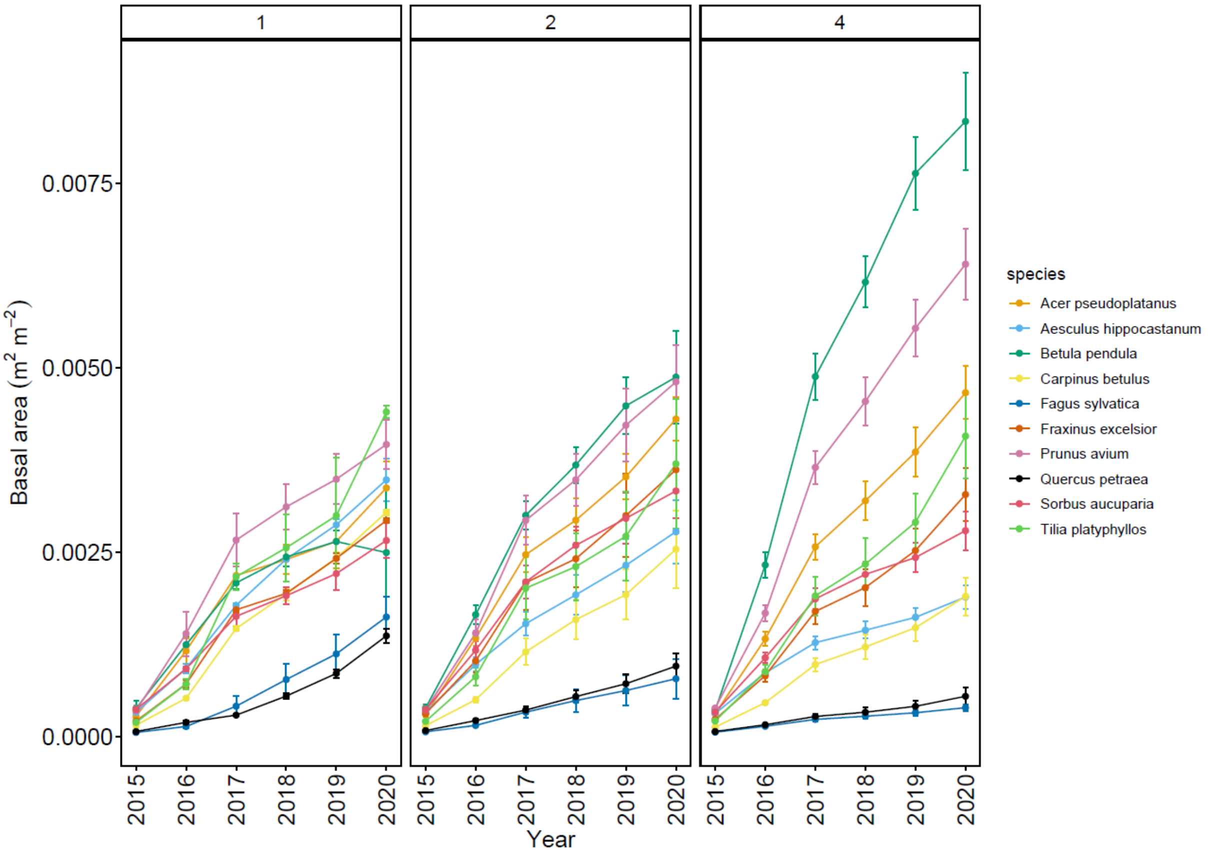


**Figure S4** Changes in basal area over time for each tree species (different colors) in 1- , 2- and 4-tree species communities. Lower number of individuals in mixtures was corrected by multiplying basal area with species richness. Dots represent mean values (± standard error) per year.

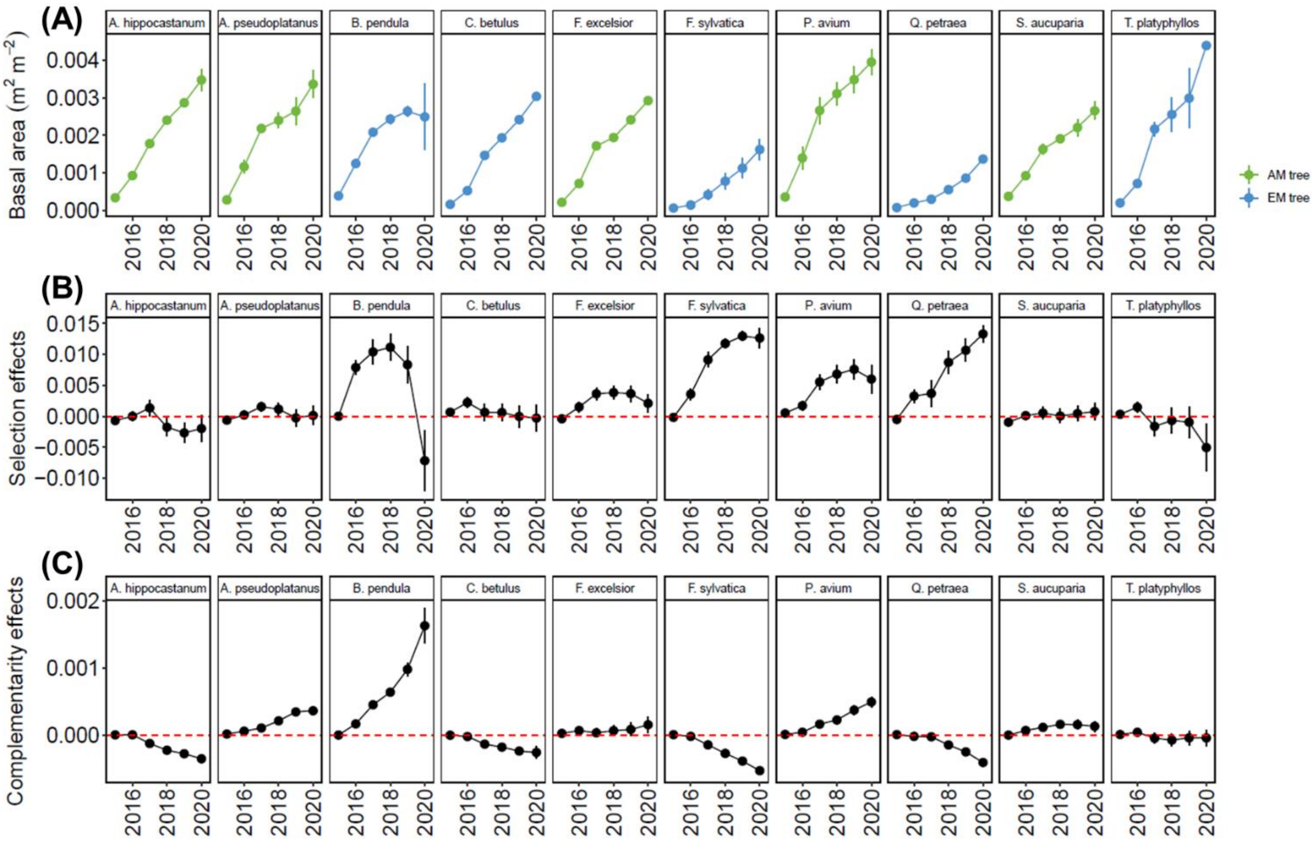


**Figure S5** Changes in basal area in monoculture (mean of both monocultures; A), species-specific selection effects (B), and species-specific complementarity effects (C) based on basal area, over time. Dots represent mean values (± standard error) per year. Color in (A) indicate whether tree species is EM or AM associated. Red dotted line indicates whether the variables are greater than zero. The y-axes of (B) and (C) are square root–transformed to reflect the quadratic nature of biodiversity effects.

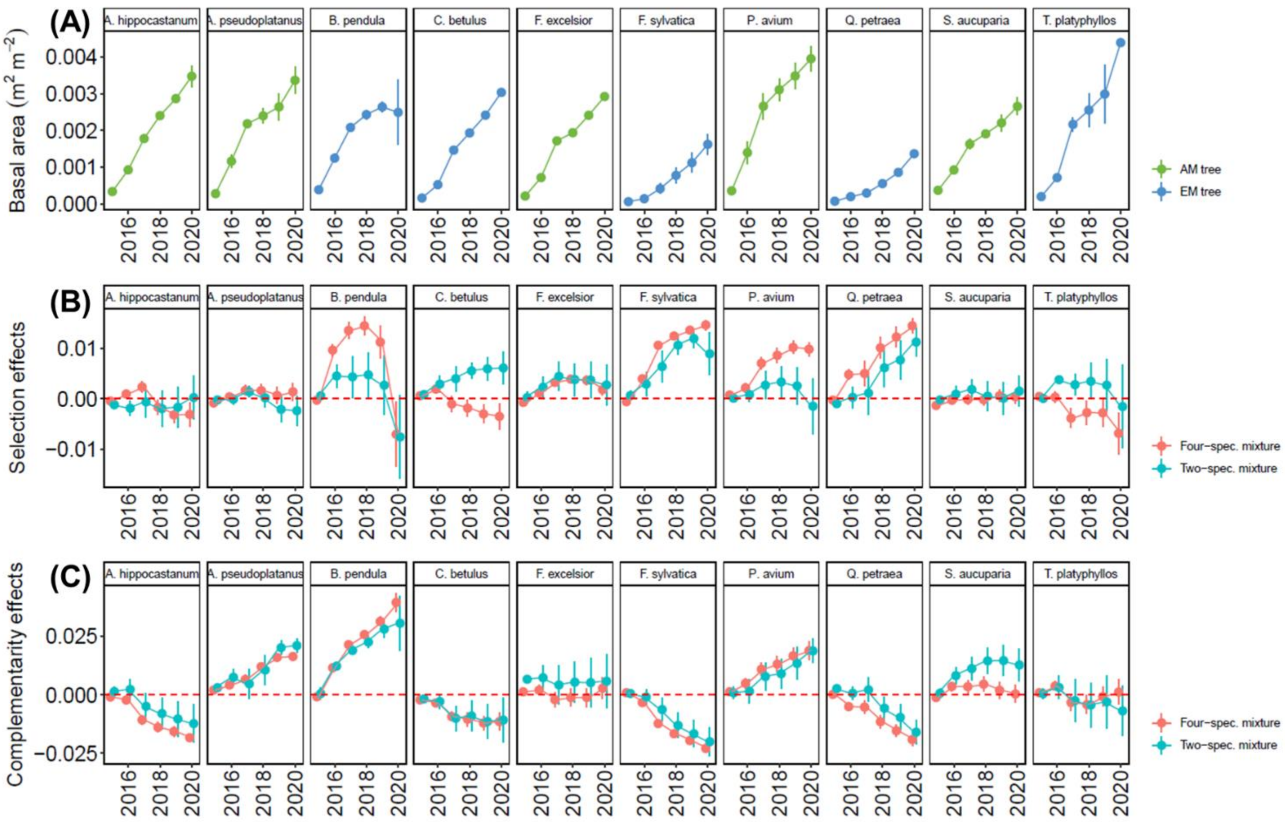


**Figure S6** Changes in basal area in monoculture (mean of both monocultures; A), species-specific selection effects (B), and species-specific complementarity effects (C) based on basal area, over time. Dots represent mean values (± standard error) per year. Colors in (A) indicate whether tree species is EM or AM associated, colors in (B) and (C) indicate effects in two- and four-tree species communities. Red dotted line indicates whether the variables are greater than zero. The y-axes of (B) and (C) are square root–transformed to reflect the quadratic nature of biodiversity effects.

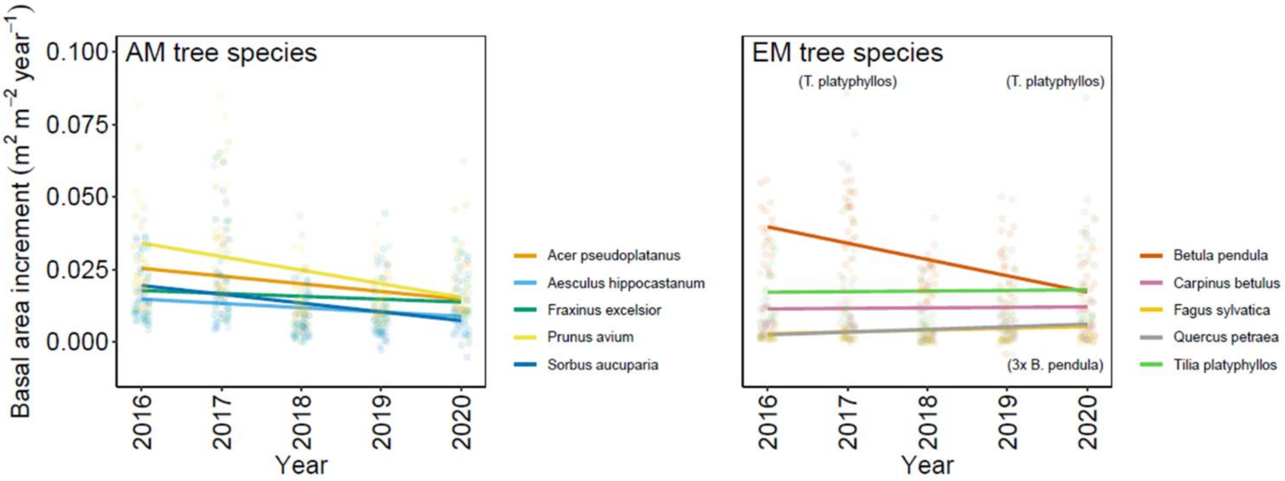


**Figure S7** Regression between basal area increment and year from 2015 to 2020 for each tree species. Each dot represents a tree community, and colors tree species. Note that five data points were excluded from the figure (for EM tree species), because they have either very high positive or negative values. These data points are indicated as text in parentheses.
