## Supplement material S2 for "Tree diversity effects on productivity depend on mycorrhizae and life strategies in a temperate forest experiment"

**Note S1** Results of community strategy analysis (basal area increment and biodiversity effects)

We found a marginally significant interaction effect of community strategy and tree species richness on basal area (and a significant three-way interaction effect (+ year) on detrended basal area). Simple linear regression analysis showed that fast and fast+slow tree communities never showed a significant relationship between species richness and basal area, while slow communities showed a positive relationship in the years 2016-2020 (Appendix S2 Table S2). For basal area increment, we found a significant interaction effect between community strategy and year (even when we corrected for differences among years). Fast and fast+slow communities decreased in increment (detrended) over the years, while slow communities increased in increment. Further, we also found a significant three-way interaction (tree species richness x community strategy x year) effect on basal area increment. This is explainable by fast+slow communities, which showed a stable increment (detrended) in two-species communities (P=0.295), while they tended to decrease in four species communities over the years (P=0.092, slope=-0.103). Furthermore, the negative slope of fast communities and the positive slope of slow communities became flatter with increasing tree species richness (fast: $slope_1$=-0.116, $slope_2$=-0.106, $slope_4$=-0.052; slow: $slope_1$=0.172, $slope_2$=0.135, $slope_4$=0.055). Results for NEs, SEs, and CEs can be found in Table S3 and Figure S3.

**Table S1** Summary of ANOVA testing differences in life/growth strategies and nutrient economy between fast and slow species. Shown are degrees of freedom (DF), F- and P-values as well as mean values for fast and slow species. Significant differences are given in bold and marginally significant differences in italics (P-values).

| | DF | F | P | Mean fast spec. | Mean slow spec. |
|---|---|---|---|---|---|
| Life/growth strategy | | | | | |
| Max. height | 1, 8 | 4.63 | *0.064* | 17.5 m | 24.0 m |
| Wood density | 1, 8 | 1.47 | 0.260 | 597 kg $m^{-3}$ | 662 kg $m^{-3}$ |
| Max. age | 1, 8 | 9.31 | **0.016** | 166 years | 350 years |
| Increment slope | 1, 8 | 11.12 | **0.010** | - 0.116 | 0.171 |
| Nutrient economy | | | | | |
| Specific leaf area | 1, 8 | 3.71 | *0.090* | 12.01 $mm^2$ $mg^{-1}$ | 17.72 $mm^2$ $mg^{-1}$ |
| Leaf CN ratio | 1, 8 | 2.18 | 0.178 | 2.3 | 1.6 |

**Table S2** Summary of simple linear regression analyses between basal area and tree species richness of fast, slow and fast+slow communities in the years 2015-2020, respectively. Shown are degrees of freedom (DF), F- and P-values as well as slope of the regression line. Significant influences are given in bold and marginally significant influences in italics (P-values).

| Basal area ~ species richness | | | | |
|---|---|---|---|---|
| Fast communities | | | | |
| | DF | F | P | slope |
| 2015 | 1 | 2.49 | 0.126 | -0.00003 |
| 2016 | 1 | 0.13 | 0.722 | -0.00002 |
| 2017 | 1 | 0.03 | 0.875 | -0.00002 |
| 2018 | 1 | 0.08 | 0.935 | <0.00001 |
| 2019 | 1 | 0.71 | 0.407 | 0.00012 |
| 2020 | 1 | 1.13 | 0.296 | 0.00019 |
| Slow communities | | | | |
| | DF | F | P | slope |
| 2015 | 1 | 2.31 | 0.139 | 0.00003 |
| 2016 | 1 | 16.98 | **<0.001** | 0.00027 |
| 2017 | 1 | 5.77 | **0.023** | 0.00040 |
| 2018 | 1 | 8.43 | **0.007** | 0.00049 |
| 2019 | 1 | 7.78 | **0.009** | 0.00055 |
| 2020 | 1 | 5.09 | **0.032** | 0.00052 |
| Fast + slow communities | | | | |
| | DF | F | P | slope |
| 2015 | 1 | 0.12 | 0.737 | -0.00002 |
| 2016 | 1 | 0.09 | 0.766 | 0.00008 |
| 2017 | 1 | 1.36 | 0.261 | 0.00045 |
| 2018 | 1 | 1.57 | 0.228 | 0.00044 |
| 2019 | 1 | 2.18 | 0.159 | 0.00062 |
| 2020 | 1 | 0.89 | 0.359 | 0.00042 |

**Table S3** Summary of mixed-effects model analyses testing the effects of log-transformed tree species richness (1, 2, 4; Sr), community strategy (fast, slow, fast+slow communitues; CS), year (2015-2020) and their interaction on net diversity effects (NEs), selection effects (SEs) and complementarity effects (CEs) based on basal area (absolute and detrended values). Shown are degrees of freedom (DF), $Chi^2$, and P-values (P). Significant influences are given in bold and marginally significant influences in italics. Note that detrended basal area values were calculated by dividing absolute basal area value per plot by the averaged value of all plots per year, to account for differences among years due to climatic fluctuations.

| | Basal area (absolute) | | | | | | |
|---|---|---|---|---|---|---|---|
| | | NEs | | SEs | | CEs | |
| | Df | $Chi^2$ | P | $Chi^2$ | P | $Chi^2$ | P |
| Tree species richness (Sr) | 1 | 2.67 | 0.102 | 6.85 | **0.009** | 0.09 | 0.768 |
| Community strategy (CS) | 2 | 8.00 | **0.018** | 14.61 | **0.001** | 0.67 | 0.715 |
| Year (Y) | 1 | 12.39 | **<0.001** | 2.29 | 0.131 | 4.16 | **0.041** |
| Sr x CS | 2 | 1.13 | 0.568 | 0.37 | 0.832 | 0.78 | 0.676 |
| Sr x Year | 1 | 12.48 | **<0.001** | 2.13 | 0.145 | 3.48 | *0.062* |
| CS x Year | 2 | 6.44 | **0.040** | 10.36 | **0.006** | 1.39 | 0.498 |
| Sr x CS x Y | 2 | 10.08 | **0.006** | 7.00 | **0.030** | 14.15 | **0.001** |
| | Basal area (detrended) | | | | | | |
| | | NEs | | SEs | | CEs | |
| | Df | $Chi^2$ | P | $Chi^2$ | P | $Chi^2$ | P |
| Tree species richness (Sr) | 1 | 1.36 | 0.244 | 6.56 | **0.010** | 0.00 | 0.960 |
| Cpmmunity strategy (CS) | 2 | 7.30 | **0.026** | 15.59 | **<0.001** | 1.05 | 0.591 |
| Year (Y) | 1 | 2.15 | 0.142 | 0.99 | 0.320 | 0.13 | 0.715 |
| Sr x CS | 2 | 0.53 | 0.768 | 0.13 | 0.937 | 0.24 | 0.887 |
| Sr x Year | 1 | 11.80 | **0.001** | 1.48 | 0.223 | 6.38 | **0.012** |
| CS x Year | 2 | 0.50 | 0.781 | 2.69 | 0.261 | 4.45 | 0.108 |
| Sr x CS x Y | 2 | 8.00 | **0.018** | 6.33 | **0.042** | 14.32 | **0.001** |

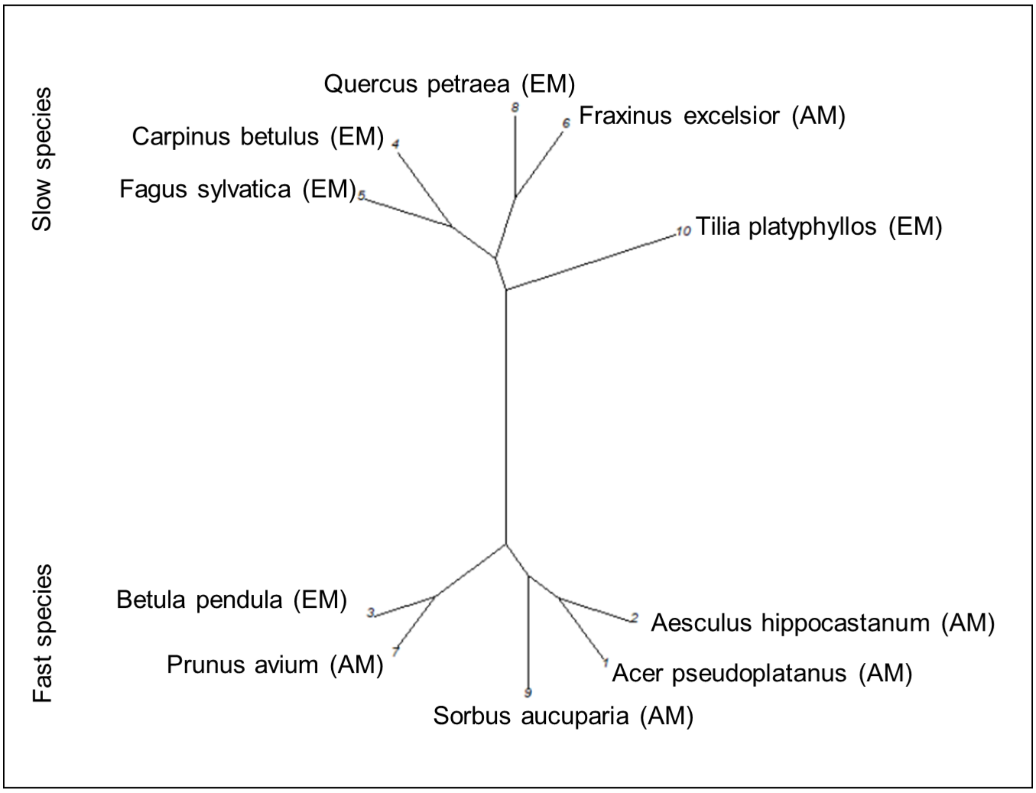


**Figure S1** Result of cluster analysis dividing the species pool into two groups (slow and fast species) based on maximum age, maximum growth height, wood density, specific leaf area, leaf C/N ratio, and increment of monoculture communities over time. Text in brackets after the species name indicate whether the species is AM or EM associated.

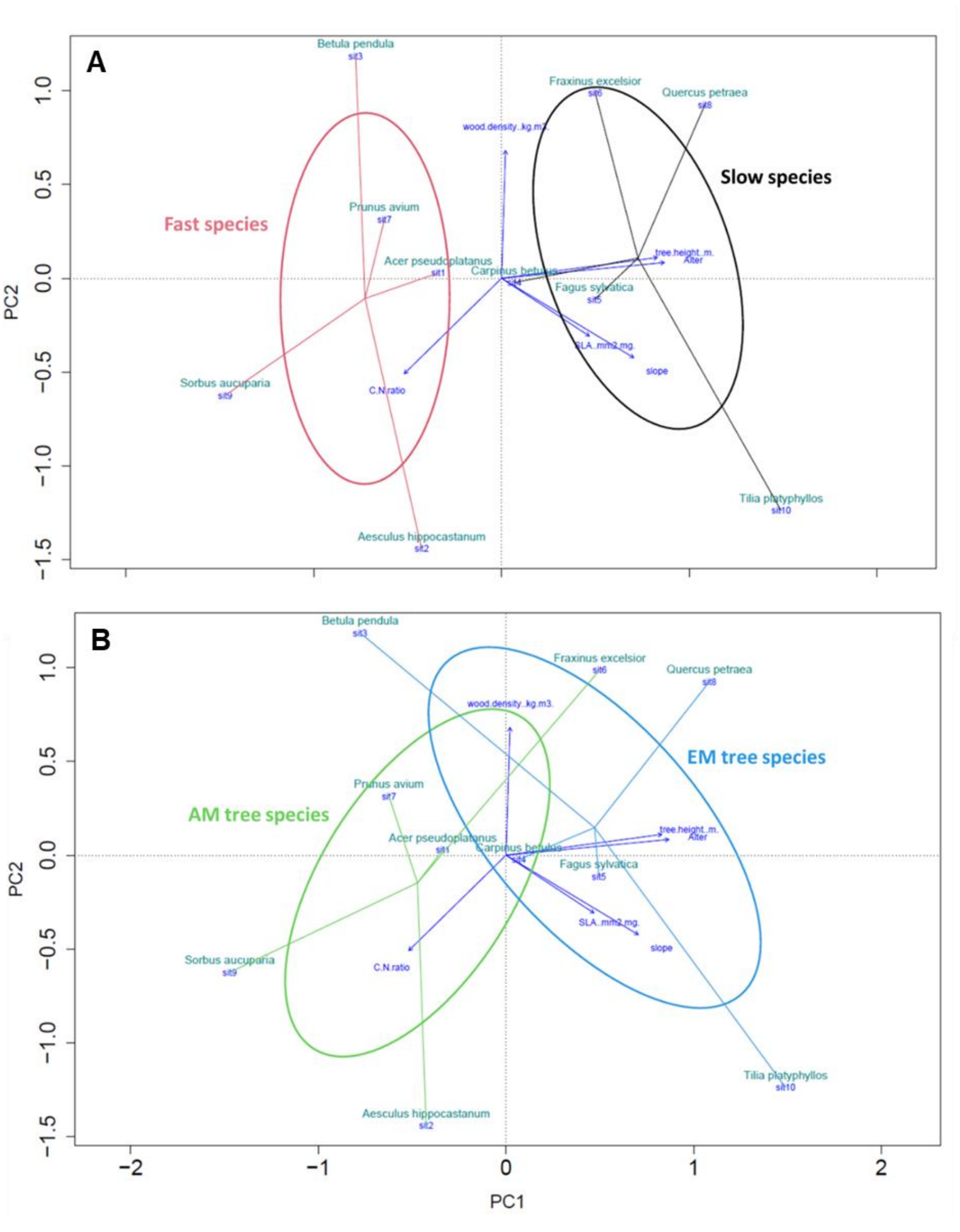


**Figure S2** Standardized principal components analyses (PCA; first versus second axes) of tree species characterized by maximum age, maximum growth height, wood density, specific leaf area, leaf C/N ratio, and increment of monoculture communities over time. Shown are life strategy groups (fast, slow; A), and mycorrhizal types (AM, EM; B) as ellipses indicating the standard deviation of point scores for each category.

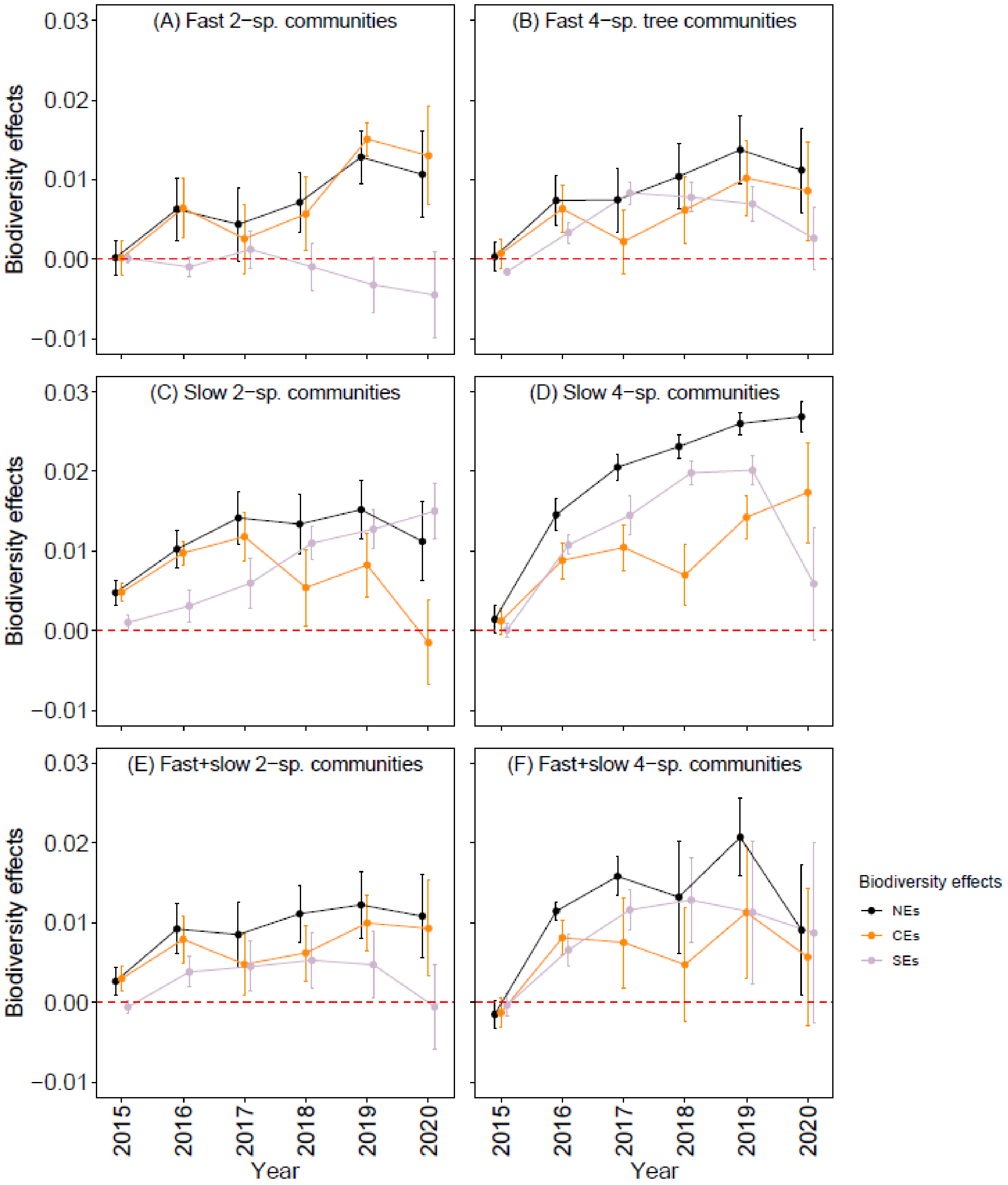


**Figure S3** Changes in net diversity effects (NEs), complementarity effects (CEs), and selection effects (SEs; different colors) over time for fast two-species (A), fast four-species (B), slow two-species (C), slow four-species (D), fast and slow two-species (E), and fast and slow four-species (F) tree communities. Dots represent mean values (± standard error) per year, and red dotted line indicates whether the biodiversity effects were greater than zero. The y-axes are square root–transformed to reflect the quadratic nature of biodiversity effects.
