## Supplement material S3 for "Tree diversity effects on productivity depend on mycorrhizae and life strategies in a temperate forest experiment"

Article title: Tree diversity effects on productivity depend on mycorrhizae and growth strategy in a temperate forest experiment

Authors: Peter Dietrich, Olga Ferlian, Yuanyuan Huang, Shan Luo, Julius Quosh, and Nico Eisenhauer

**Note S1** Calculation of biomass production and statistical analysis

For calculation of stem biomass production per tree and year, we used basal diameter ($d_0$; m) and height (m), and species-specific allometric equations, which can be found in Kaulen et al. (in prep.). Further, calculations of tree community biomass production, biomass increment, and statistical analyses of these variables were performed in the same way as discussed in the Materials and Methods section for basal area.

**Table S1** Summary of mixed-effects model analyses testing the effects of log-transformed tree species richness (1, 2, 4; Sr), mycorrhizal type (AM, EM, AM+EM; Myc), year (2015-2020) and their interaction on log-transformed biomass and non-transformed biomass values (absolute and detrended values). Shown are degrees of freedom (DF), $Chi^2$, and P-values (P). Significant influences are given in bold and marginally significant influences in italics. Note that detrended values were calculated by dividing absolute value per plot by the averaged value of all plots per year, to account for differences among years due to climatic fluctuations.

| | | Biomass (log) | | | | Biomass (non-transformed) | | | |
|---|---|---|---|---|---|---|---|---|---|
| | | absolute | | detrended | | absolute | | detrended | |
| | Df | $Chi^2$ | P | $Chi^2$ | P | $Chi^2$ | P | $Chi^2$ | P |
| Tree species richness (Sr) | 1 | 6.50 | **0.011** | 6.50 | **0.011** | 0.14 | 0.705 | 0.46 | 0.497 |
| Mycorrhizal type (Myc) | 2 | 7.00 | **0.030** | 7.00 | **0.030** | 0.39 | 0.825 | 2.16 | 0.339 |
| Year | 1 | 22.26 | **<0.001** | 1.74 | 0.188 | 48.41 | **<0.001** | 0.00 | 1.000 |
| Sr x Myc | 2 | 1.76 | 0.415 | 1.76 | 0.416 | 0.18 | 0.916 | 0.19 | 0.912 |
| Sr x Year | 1 | 2.43 | 0.119 | 2.47 | 0.116 | 0.01 | 0.914 | 0.20 | 0.654 |
| Myc x Year | 2 | 84.00 | **<0.001** | 85.31 | **<0.001** | 10.03 | **0.007** | 50.21 | **<0.001** |
| Sr x Myc x Year | 2 | 5.45 | *0.066* | 5.53 | *0.063* | 1.16 | 0.561 | 0.74 | 0.691 |

**Table S2** Summary of mixed-effects model analyses testing the effects of log-transformed tree species richness (1, 2, 4; Sr), mycorrhizal type (AM, EM, AM+EM; Myc), year (2016-2020) and their interaction on biomass increment (absolute and detrended values). Shown are degrees of freedom (DF), Chi$^2$, and P-values (P). Significant influences are given in bold and marginally significant influences in italics. Note that detrended values were calculated by dividing absolute value per plot by the averaged value of all plots per year, to account for differences among years due to climatic fluctuations.

| | | Biomass increment | | | |
|---|---|---|---|---|---|
| | | absolute | | detrended | |
| | Df | Chi$^2$ | P | Chi$^2$ | P |
| Tree species richness (Sr) | 1 | 0.01 | 0.915 | <0.01 | 1.000 |
| Mycorrhizal type (Myc) | 2 | 2.81 | 0.245 | 2.43 | 0.296 |
| Year | 1 | 2.10 | 0.148 | <0.01 | 1.000 |
| Sr x Myc | 2 | 0.21 | 0.900 | 0.19 | 0.911 |
| Sr x Year | 1 | 4.72 | **0.030** | 6.11 | **0.013** |
| Myc x Year | 2 | 30.17 | **<0.001** | 32.92 | **<0.001** |
| Sr x Myc x Year | 2 | 0.91 | 0.634 | 1.22 | 0.542 |

**Table S3** Summary of mixed-effects model analyses testing the effects of log-transformed tree species richness (1, 2, 4; Sr), mycorrhizal type (AM, EM, AM+EM; Myc), year (2015-2020) and their interaction on net diversity effects (NEs), selection effects (SEs) and complementarity effects (CEs) based on biomass (absolute values). Shown are degrees of freedom (DF), Chi$^2$, and P-values (P). Significant influences are given in bold and marginally significant influences in italics.

| | | Biomass (absolute) | | | | | |
|---|---|---|---|---|---|---|---|
| | | NEs | | SEs | | CEs | |
| | Df | Chi$^2$ | P | Chi$^2$ | P | Chi$^2$ | P |
| Tree species richness (Sr) | 1 | 0.29 | 0.588 | 0.14 | 0.708 | <0.01 | 0.964 |
| Mycorrhiza type (Myc) | 2 | 3.54 | 0.170 | 5.15 | *0.076* | 2.28 | 0.320 |
| Year | 1 | 0.50 | 0.480 | 0.49 | 0.484 | 2.29 | 0.131 |
| Sr x Myc | 2 | 0.18 | 0.915 | 0.15 | 0.926 | 0.61 | 0.737 |
| Sr x Year | 1 | 3.33 | *0.068* | 9.50 | **0.002** | 2.38 | 0.123 |
| Myc x Year | 2 | 19.77 | **<0.001** | 22.90 | **<0.001** | 1.86 | 0.395 |
| Sr x Myc x Year | 2 | 3.59 | 0.166 | 4.62 | *0.099* | 3.35 | 0.187 |

**Table S4** Summary of mixed-effects model analyses testing the effects of log-transformed tree species richness (1, 2, 4; Sr), community strategy (fast, slow, fast+slow communities; CS), year (2015-2020) and their interaction on log-transformed and non-transformed biomass values (absolute and detrended values). Shown are degrees of freedom (DF), $Chi^2$, and P-values (P). Significant influences are given in bold and marginally significant influences in italics. Note that detrended values were calculated by dividing absolute value per plot by the averaged value of all plots per year, to account for differences among years due to climatic fluctuations.

| | | Biomass (log) | | | | Biomass (non-transformed) | | | |
|---|---|---|---|---|---|---|---|---|---|
| | | absolute | | detrended | | absolute | | detrended | |
| | Df | $Chi^2$ | P | $Chi^2$ | P | $Chi^2$ | P | $Chi^2$ | P |
| Tree species richness (Sr) | 1 | 6.50 | **0.011** | 6.50 | **0.011** | 0.14 | 0.705 | 0.46 | 0.497 |
| Community strategy (CS) | 2 | 7.00 | **0.030** | 7.00 | **0.030** | 0.39 | 0.825 | 2.16 | 0.339 |
| Year | 1 | 22.26 | **<0.001** | 1.74 | 0.188 | 48.41 | **<0.001** | 0.00 | 1.000 |
| Sr x CS | 2 | 1.76 | 0.415 | 1.76 | 0.416 | 0.18 | 0.916 | 0.19 | 0.912 |
| Sr x Year | 1 | 2.43 | 0.119 | 2.47 | 0.116 | 0.01 | 0.914 | 0.20 | 0.654 |
| LS x Year | 2 | 84.00 | **<0.001** | 85.31 | **<0.001** | 10.03 | **0.007** | 50.21 | **<0.001** |
| Sr x CS x Year | 2 | 5.45 | *0.066* | 5.53 | *0.063* | 1.16 | 0.561 | 0.74 | 0.691 |

**Table S5** Summary of mixed-effects model analyses testing the effects of log-transformed tree species richness (1, 2, 4; Sr), community strategy (fast, slow, fast+slow communities; CS), year (2015-2020) and their interaction on annual biomass increment (absolute and detrended values). Shown are degrees of freedom (DF), $Chi^2$, and P-values (P). Significant influences are given in bold and marginally significant influences in italics. Note that detrended values were calculated by dividing absolute value per plot by the averaged value of all plots per year, to account for differences among years due to climatic fluctuations.

| | | Biomass increment | | | |
|---|---|---|---|---|---|
| | | absolute | | detrended | |
| | Df | $Chi^2$ | P | $Chi^2$ | P |
| Tree species richness (Sr) | 1 | 0.01 | 0.915 | <0.01 | 1.000 |
| Community strategy (CS) | 2 | 1.09 | 0.579 | 0.55 | 0.761 |
| Year | 1 | 2.09 | 0.148 | <0.01 | 1.000 |
| Sr x CS | 2 | 0.25 | 0.884 | 0.15 | 0.927 |
| Sr x Year | 1 | 4.72 | **0.030** | 6.11 | **0.013** |
| CS x Year | 2 | 50.50 | **<0.001** | 65.21 | **<0.001** |
| Sr x CS x Year | 2 | 8.40 | **0.015** | 11.99 | **0.002** |
